# Computational analysis of threonine ladders on distinct beta-solenoid scaffolds, with implications for the design of novel antifreeze proteins

**DOI:** 10.1101/2025.03.18.644030

**Authors:** Cianna N. Calia, Francesco Paesani

## Abstract

Cold-adapted organisms frequently express antifreeze proteins (AFPs) that facilitate their survival at low temperatures, with some especially potent insect AFPs exhibiting beta-solenoid structures with ice-binding threonine ladders. Beta-solenoids exist in nature in numerous forms and emerging protein design technologies may afford opportunities to diversify them further, suggesting the possibility of developing a variety of new AFPs by installing a threonine ladder on non-AFP natural or designed beta-solenoids. However, early attempts at such engineering, combined with differences observed between AFPs and structurally similar ice-nucleating proteins, have raised a critical question: Will a threonine ladder show essentially the same behavior regardless of the beta-solenoid scaffold that hosts it, or does the specific solenoid scaffold significantly affect a threonine ladder’s structural characteristics (and thus potentially alter its suitability for ice binding)? We set out to address this question by creating distinct variants of a simplified model beta-solenoid for *in silico* analysis via structure prediction and molecular dynamics simulations. Our findings indicate that local structural details such as the distance between the hydroxyl groups of adjacent threonines in a TXT motif can vary depending on the beta-solenoid scaffold. In the most extreme example among our model solenoids, we observed in simulations that differences in only inward-facing residues of the scaffold were sufficient to influence the presence of ordered channel waters between the threonines, a noted feature of natural ice-binding threonine surfaces such as that of TmAFP. While additional studies will be necessary to expand on how such distinctions affect activity, these results emphasize that the impact of a particular beta-solenoid scaffold on the local geometry of a threonine ladder may be a pertinent consideration in future efforts to design novel hyperactive AFPs to support applications ranging from biomedical cryopreservation to food science. We conclude our present investigation with a preliminary exploration of how this and other considerations manifest in a proposed workflow for generating predicted AFP-like beta-solenoids using AlphaFold and ProteinMPNN.

## Introduction

Antifreeze proteins (AFPs) are a key survival mechanism for diverse organisms adapted to icy environments, from psychrophilic microbes to polar fish to overwintering insects and plants.^1–4^ AFPs are a class of ice-binding proteins (IBPs) that exert thermal hysteresis activity via the Kelvin effect: They lower the freezing temperature by binding to ice crystals and inducing curvature that raises the free energy of the ice phase. ^5–7^ They also exhibit ice recrystallization inhibition, impeding the formation of large ice crystals at the expense of smaller ones.^8,9^ These forms of activity make AFPs a promising technological solution for applications such as cryopreservation of cells, tissues, and organs for research and medical use;^10–15^ protection of crops from frost damage;^16–19^ and texture modulation in foods such as ice cream.^20–23^

AFPs exhibit a range of structures, such as a single alpha helix in the type I AFP from winter flounder,^24^ a small globular fold in the type III AFP from ocean pout,^25^ and beta-solenoid structures in hyperactive insect AFPs.^26,27^ All known AFPs are believed to possess an ice-binding surface (IBS), though the nature of the IBS varies with AFP type. Hyperactive insect AFPs such as TmAFP (from the mealworm beetle *Tenebrio molitor*) and sbwAFP (from the spruce budworm *Choristoneura fumiferana*), which have especially potent thermal hysteresis activity,^28,29^ have aligned threonine-X-threonine (TXT) motifs in the consecutive loops of their beta-solenoid structures, forming a flat face with two rows of outward-facing threonine residues — a threonine ladder IBS ^26,27,30^ (Figure 1A). Numerous studies have identified ordered water molecules, such as channel waters between the threonine rows, as an additional and critical feature of the IBS (e.g., Refs. 26,31–41).

**Figure 1:**
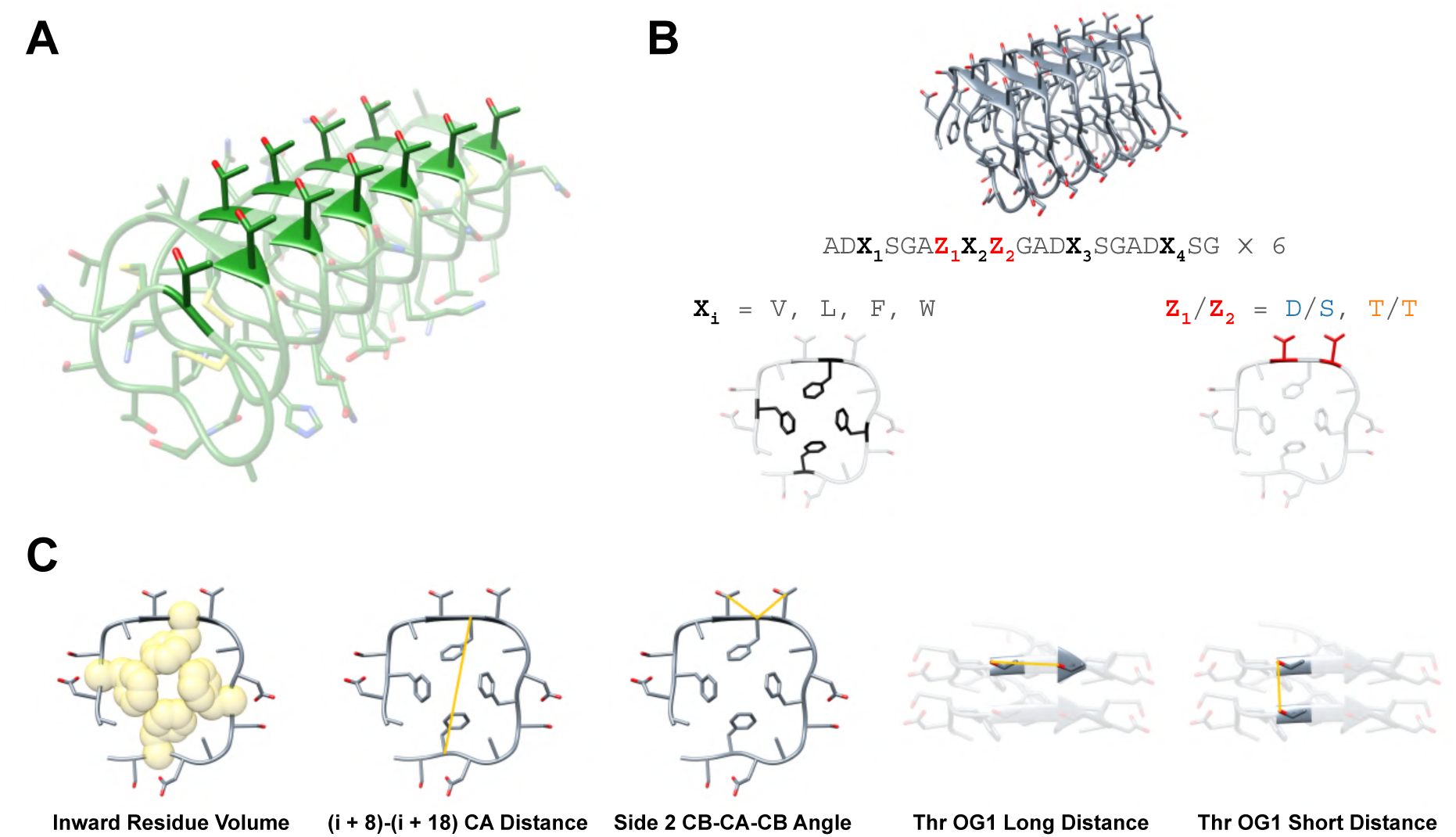
(A) Crystal structure of TmAFP (PDB 1EZG), with IBS threonines highlighted in darker green. (B) Overview of our construction of model beta-solenoid sequences based on pentapeptide repeat proteins. (C) Visual outline of the key features relevant to our analysis of predicted structures and simulations.

The relative simplicity of the TXT motif and the high thermal hysteresis activity of insect AFPs together suggest that the threonine ladder may be an ideal IBS for use in developing novel AFPs to broaden the set of ice-binding biomolecules available for various applications. For instance, transplanting the threonine ladder IBS onto alternative beta-solenoid scaffolds that are not stabilized by disulfide bonds presents one potential strategy to create highly active new AFPs with reduced expression challenges compared to insect AFPs that rely on extensive disulfide bonding for stability.^42^ With the recent explosion of deep learning methods for protein design and structure prediction (e.g., Refs. 43–53), even more creative possibilities for AFP/IBP design and optimization are now also within reach. However, whether a threonine ladder can function equally well as an IBS on distinct scaffolds, either natural or designed, that did not evolve for this role remains an open question.

Pentapeptide repeat proteins are a class of bacterial proteins that adopt right-handed beta-solenoid structures with a square profile. Each square loop contains four sets of five residues; the inward-facing residue in the middle of each side is typically leucine or phenylalanine.^54,55^ Importantly, pentapeptide repeat proteins have relatively flat faces, do not require disulfide bonding for stabilization, and tolerate a variety of different residues in their outward-facing positions.^56^ These properties make pentapeptide repeat proteins a promising choice as new beta-solenoid AFP scaffolds. Indeed, Yu introduced a threonine ladder into a protein from this class, and the mutant displayed ice shaping and thermal hysteresis activity.^57^ This novel AFP’s thermal hysteresis activity was about an order of magnitude less than that of TmAFP, however, highlighting the possibility that some beta-solenoids may be more optimal AFP scaffolds than others.

This possibility is further supported by comparisons between AFPs and bacterial ice-nucleating proteins (INPs), which also have a beta-solenoid structure displaying a threonine ladder.^58–60^ Recent computational studies have suggested that the ice-binding threonine ladder on sbwAFP differs from that on an INP segment in terms of local water structure and dynamics^40^ as well as ice nucleating ability. ^61^ Furthermore, like Yu’s design, a truncated INP showed substantially reduced freezing point depression compared to hyperactive AFPs.^62^ Plausible reasons for disparities in thermal hysteresis activity among different beta-solenoids with ice-binding TXT sequence motifs include reduced binding due to altered threonine geometry, suboptimal scaffold shape or non-binding surface chemistry for resisting engulfment by ice, the amount of twist in the solenoid, cooperative effects or additive structural mismatch dependent on the number of solenoid loops, the presence of any intervening non-threonine residues in the IBS, and variations in solenoid rigidity.

Here, we devise simplified comparisons to investigate the first of those considerations: whether and how a threonine ladder’s structural characteristics depend on the nature of the beta-solenoid scaffold. Liou et al. highlighted the remarkable regularity of the crystal structure of TmAFP, with its threonine hydroxyl groups exhibiting spacings nearly identical to those of water oxygens in the ice lattice. ^26^ We hypothesized that threonine ladders on distinct solenoid scaffolds may show differences in local structure that would potentially render them less optimal for ice binding, and that quantifying these differences may help inform future efforts to design novel beta-solenoid AFPs. Using repetitive sequence patterns based on pentapeptide repeat proteins as a convenient model system, we employed a combination of structure predictions and molecular dynamics (MD) simulations to assess how changes in the beta-solenoid scaffold impact the threonine ladder structure, as quantified by simple geometric features for a set of hypothetical *in silico* variants. Together these analysis methods demonstrated the scaffold’s ability to influence the threonine ladder, with less concave solenoid faces providing the closest approximation to the IBS of TmAFP and with the hydroxyl oxygen spacings between adjacent threonines within the same TXT motif showing more noticeable variation than those between aligned threonines in consecutive loops. Finally, we explore possible practical implications of our findings in a protein design context by examining preliminary AFP-like beta-solenoid designs among sequences generated using the deep learning tool ProteinMPNN.^46^

## Methods

### Model beta-solenoid variants

An initial model solenoid loop sequence was constructed by repeating the pentapeptide repeat consensus motif ADLSG^54,56^ four times. The inward-facing residue (X) at the center of each of the four sides of the 20-residue loop was substituted with V, F, or W or kept as L to create a non-exhaustive set of 40 distinct loops (Figure 1B). Each loop was repeated six times to construct the threonine-free variants (labeled X_1_X_2_X_3_X_4_ DS). A threonine ladder was introduced into each variant by replacing the DXS on one face (side 2 counting from the N-terminus) with TXT in every loop, yielding 40 additional variants of the form X_1_X_2_X_3_X_4_ TT. The 20-residue loop sequences for all variants are listed in Tables S1 and S2 of the Supporting Information. The average inward residue volume for a given sequence refers to the average volume of residues X_1_, X_2_, X_3_, and X_4_ based on the consensus volumes reported in Ref. 63.

### Structure prediction and analysis

Except where otherwise indicated, our AlphaFold predictions refer to results from running ColabFold v1.5.5^64^ in single-sequence mode with three recycles, without templates, and using the AlphaFold 2 ptm weights (all five models). ^43^ The structures were not relaxed. These settings were sufficient to obtain many highly confident solenoid predictions (Figures S2A and S2B of the Supporting Information). We further reasoned that omitting multiple sequence alignments and templates may help avoid biasing our analysis toward sequences more similar to those found in nature, as has previously been noted in the context of protein design workflows.^48^ Structure ranks refer to ranking each set of five structures by average pLDDT. Our AlphaFold 3^49^ comparisons were run on AlphaFold Server with all default settings; mmCIF files from the server were converted to PDB format with the GEMMI web-based converter.^65^

We employed Biopython^66^ and PYTRAJ,^67^ the Python interface for CPPTRAJ,^68^ for structure analysis. Note that we use atom names consistent with the Amber force field (see below) throughout our results (Figure S1 of the Supporting Information) and we refer to features by the names shown in Figure 1C. Angle and distance values for TmAFP, sbwAFP501, and sbwAFP337 were taken from PDB 1EZG,^26^ 1M8N,^31^ and 1L0S,^69^ respectively. Threonines in predicted structures were considered to be in the intended rotameric state if their N-CA-CB-OG1 dihedral angle was ≥ –60*^◦^* and ≤ –50*^◦^*; the upper bound was changed to –45*^◦^* for our ProteinMPNN sequences to account for slight scaffold-dependent differences.

### MD simulations and analysis

All simulations were run with the CUDA-accelerated implementation of PMEMD in Amber22.^70–74^ Chain A of PDB 1EZG (for TmAFP) and the top-ranked AlphaFold 2 structures for the LLLL TT, FFFF TT, FFFF DS, and WWWF TT pentapeptide repeat variants and for the selected ProteinMPNN sequences were each solvated in a cubic water box using tLEaP from AmberTools,^70^ with a 15 Å minimum buffer distance. The rank 3 structure of LLLL TT was also used. The ff14SB force field^75^ was employed for the proteins, and water was represented by the TIP4P-Ew model.^76^ These parameters were sufficient to reproduce expected details of TmAFP’s IBS such as the dominant rotameric state of the threonines^36,77,78^ (Figures S3A and S3B of the Supporting Information) and the presence of channel waters^26,35,36,38,41^ (Figure 5E).

Each system underwent an energy minimization followed by heating in the *NV T* (constant number of particles, volume, and temperature) ensemble over 50 ps to reach the desired temperature (300 K or 242 K), with harmonic backbone restraints applied during both stages using a force constant of 10 kcal/mol·Å^2^. Next, for each system and each temperature we ran a 10-ns equilibration in the *NPT* (constant number of particles, pressure, and temperature) ensemble at 1 bar with weakened backbone restraints (2 kcal/mol·Å^2^). We then performed unbiased 100-ns *NPT* production simulations with no restraints, saving frames every 10 ps. Throughout the simulations we employed periodic boundary conditions, a cutoff of 12 Å for non-bonded interactions, a timestep of 2 fs, the SHAKE algorithm^79^ to constrain bonds involving hydrogen atoms, the Langevin thermostat, and the Berendsen barostat (in *NPT* simulations). Example Amber inputs are included in the Supporting Information.

We analyzed simulation trajectories using CPPTRAJ. For rotamer-filtered threonine OG1 distance distributions, only frames in which both threonines had the N-CA-CB-OG1 dihedral angle ≥ –75*^◦^* and ≤ –50*^◦^* were considered; these cutoffs were selected based on Figures S3A and S3B of the Supporting Information. For 4MZU sample1, the limits were changed to –70*^◦^* and –45*^◦^* based on Figure S13A of the Supporting Information. We computed the three-dimensional water oxygen density distributions in a rectangular space around the threonine ladders in the low-temperature simulations with CPPTRAJ’s grid command using a grid spacing of 0.5 Å.

### ProteinMPNN

Except for our tests of different sampling temperatures for the LLLL TT structure, we ran ProteinMPNN with a temperature of 0.8 in order to promote sequence diversity (see Figure S10 of the Supporting Information). We used the v 48 020 model, excluded cysteines, and fixed the two rows of threonines on the selected TXT face for each case.

### Structure and data visualizations

All structure visualizations were created with UCSF Chimera,^80^ including images of water oxygen density distributions produced with Chimera’s Volume Viewer tool. Python and Matplotlib^81^ were used for all plots; distribution plots were made with the Seaborn library^82^ using kernel density estimation. Sequence logos were generated with WebLogo.^83^

## Results

### Structure predictions for variations of a model beta-solenoid

We began by constructing a highly regular model beta-solenoid by repeating the pentapeptide repeat consensus pattern ADLSG 24 times to form six four-sided loops. From this starting sequence, we diversified the solenoid scaffold by creating 40 variants differing in the volume of the inward-facing hydrophobic residues located at the center of each face of the repeated square loop. For each of these variants we made a version with the outward-facing residues on one side changed to threonines, converting the DXS pattern to a TXT motif in each loop. The hydrophobic residues placed in the central positions (L, V, F, or W) encompass variations both within and beyond the natural sequence space of pentapeptide repeat proteins^54^ and are not necessarily intended to yield realistic proteins, but rather to provide a simple strategy for placing threonine ladders in subtly distinct structural contexts. Each variant is henceforth referred to as X_1_X_2_X_3_X_4_ Z_1_Z_2_, where X*_i_* is the central inward-facing residue on side *i* of the repeated loop and Z_1_/Z_2_ are the outward-facing residues on side 2 (the second side counting from the N-terminus) (Figure 1B). According to this system, the original consensus-based sequence consisting of 24 repeats of ADLSG is labeled LLLL DS. We used the ColabFold implementation of AlphaFold 2 to predict structures for all 80 resulting sequences (40 with a threonine ladder and 40 without, with loop sequences listed in Tables S1 and S2 of the Supporting Information).

The majority of the predicted structures had high pLDDT values and low alpha-carbon RMSD with respect to the highest-ranked predicted structure of LLLL DS, suggesting that AlphaFold readily predicts this structure regardless of deviations from natural sequences in the pentapeptide repeat protein family. Lower pLDDT and greater structural variation occurred in a subset of cases involving particularly high or low inward residue volumes (mainly tryptophan or valine) (Figures S2A and S2B of the Supporting Information). Only highly confident beta-solenoid structures were considered for further analysis, with average pLDDT greater than 90 and CA RMSD to LLLL DS (rank 1) less than 2.0 Å. Out of the 247 predicted structures meeting those criteria, nine exhibited topological deviations in which the corner alanines were flipped outward (Figure S2C of the Supporting Information), disrupting the typical inward-outward-inward-outward-outward pattern; these cases were also excluded from subsequent analysis.

We emphasize that even the confidently predicted solenoids among this set are intended only as *in silico* models for assessing how the threonines may be influenced by scaffold changes; these are useful analysis tools, not necessarily promising designed proteins. Many of these structures, particularly those containing unnatural tryptophan or valine substitutions, may be fictitious, consistent with a recent description of AlphaFold 2’s propensity to hallucinate beta-solenoids.^84^ AlphaFold’s tendency to produce this structure even when unrealistic inward-facing residues are present may limit its predictive ability as a tool for screening designed solenoid sequences, but on the other hand made this fold convenient for our present analysis because we could systematically introduce a range of relatively subtle scaffold differences based on the sizes of the core hydrophobes.

Treating the average volume of the central inward-facing residues as the independent variable, we defined a set of simple geometric features that could readily be calculated from the predicted structures to assess the impact of differences in the inward residues. These features are illustrated in Figure 1C and are denoted using atom names depicted in Figure S1 of the Supporting Information. Terminal loops were not included in the calculation of these features because we expected greater structural variability near the termini.

As AlphaFold can sometimes be robust to minor sequence changes and thus imperfect for predicting the effects of mutations,^85,86^ we first sought to confirm that our AlphaFold structures showed a physically reasonable response to the differences in inward residue bulk, i.e., that we had indeed introduced variation in the scaffold. We reasoned that the square beta-solenoid profile may progressively bulge as the inward residues increase in size. Consistent with this expectation, the distance between the alpha carbon atoms of the eigth and 18th residues in each loop (the central inward residues on sides 2 and 4) generally increased with average inward residue volume, both in the variants with threonine ladders and in those without (Figure 2A). This feature, henceforth referred to as the (i + 8)-(i + 18) CA distance (with i representing the number of residues before the start of the loop in question), should not be directly relevant for ice binding but demonstrated AlphaFold’s ability to show differences in beta-solenoids with identical topologies.

**Figure 2:**
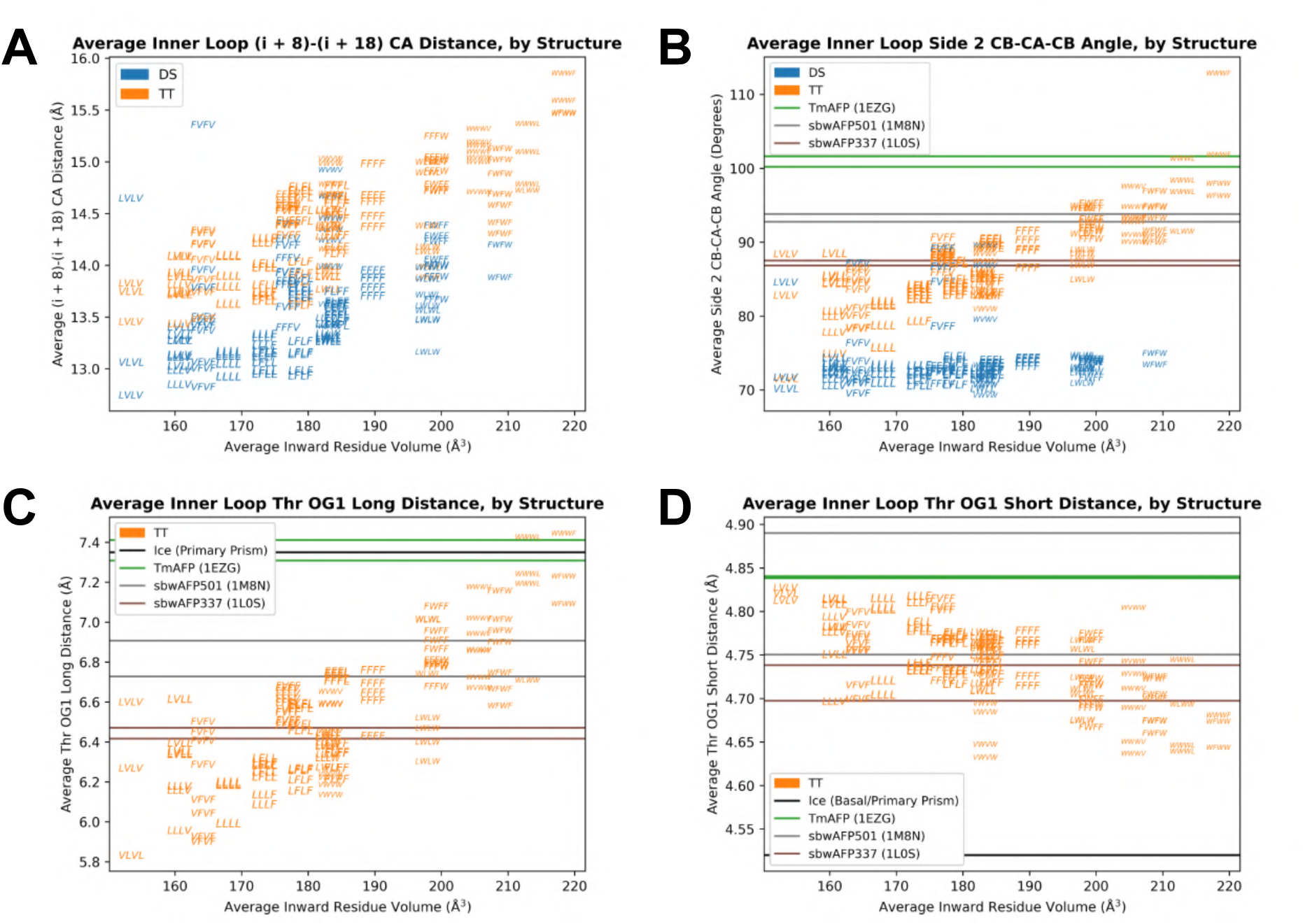
Geometric features from AlphaFold structures with respect to the average inward residue volume for the pentapeptide repeat variants, including (A) the (i + 8)-(i + 18) CA distance, (B) the side 2 CB-CA-CB angle, (C) the threonine OG1 long distance, and (D) the threonine OG1 short distance. Each data point, representing an average over the four inner (non-terminal) loops in one of the five predicted structures for a given sequence, is denoted by the inward residues of the variant’s four faces (X_1_X_2_X_3_X_4_), with the Z_1_/Z_2_ residues indicated by the color (blue for threonine-free variants and orange for those with threonine ladders). Only structures with average pLDDT greater than 90 and CA RMSD to LLLL DS rank 1 less than 2.0 Å and without topological deviations are included in (A) through (D), and only structures which also had all inner loop threonines in the expected rotameric state are included in (C) and (D). In (B) through (D) values from TmAFP and sbwAFP are shown by horizontal lines for comparison (chains A and B of PDB 1EZG, 1M8N, and 1L0S), along with oxygen spacings from ice^26^ in (C) and (D).

Next we assessed structural features specific to the TXT face (or the analogous DXS face in the threonine-free variants). We supposed that the inward residues might influence the concavity of the beta-solenoid’s faces, as quantified for our selected face by the side 2 CB-CA-CB angle, which could in turn affect the spacings between the threonines’ hydroxyl oxygen (OG1) atoms. For the variants with threonine ladders, the AlphaFold structures showed the trend we expected: The side 2 CB-CA-CB angle increased with inward residue volume, indicating that the threonine ladder face became less concave (Figure 2B). However, this trend was absent for the corresponding DXS face of the threonine-free variants. We suspect that hydrogen bonding involving the aspartate and serine side chains might make the geometry of the DXS motif relatively indifferent to the inward residues, unlike threonines which appeared more sensitive to changes in the scaffold.

Furthermore, the distance between the hydroxyl oxygens of adjacent threonines in the same loop (henceforth called the “OG1 long distance”) showed a positive correlation with inward residue volume (Figure 2C). The “OG1 short distance” (between adjacent threonine oxygens in consecutive loops) showed a negative correlation but over a much narrower range of values (Figure 2D), entirely within the range between the data points from TmAFP’s crystal structure and the ice lattice spacing of 4.52 Å.^26^ Only structures with threonines in the same rotameric state as those in TmAFP’s crystal structure were considered in Figures 2C and 2D; most structures met this criterion, exhibiting N-CA-CB-OG1 dihedral angles between –60*^◦^* and –50*^◦^* (Figure S2D of the Supporting Information).

For both the side 2 angle and the OG1 long distance (Figures 2B and 2C), the bulkiest inward residues yielded values closest to those from the crystal structure of TmAFP, which exhibited OG1 long distances much nearer to the longer oxygen spacings in the primary prism (7.35 Å) and basal (7.83 Å) planes^26^ of ice than did the model solenoids at the low end of the inward residue volume distribution. The side 2 angles and OG1 long distances from the crystal structures of two different isoforms of sbwAFP fell within the intermediate parts of these ranges, with sbwAFP501 nearer to TmAFP. sbwAFP337 was farther from TmAFP but still exhibited larger side 2 angle and OG1 long distance values than the model structures containing the consensus-based inward-facing leucines. We note that TmAFP constitutes the most direct comparison to our pentapeptide repeat variants in that it is a right-handed beta-solenoid with a continuous array of eight threonines (four TXT motifs) in its interior loops; furthermore it crystallized with channel waters present,^26^ consistent with simulations of the protein in solution.^35–38,41^ Comparisons to sbwAFP may involve additional considerations: The sbwAFP isoforms are left handed, neither isoform contains an uninterrupted stretch of four non-terminal TXT loops, and sbwAFP337 crystallized in a dimeric arrangement that precluded the presence of channel waters that are nevertheless believed to exist when the monomer is in solution.^69^

### Assessment of structural trends with MD simulations

To determine whether the trends from AlphaFold’s static structure predictions would be upheld by physics-based methods that incorporate dynamics, we selected four variants for analysis with MD simulations under ambient conditions. We chose the variants LLLL TT, FFFF TT, and WWWF TT to encompass cases with low, intermediate, and high inward residue volume, respectively, as well as FFFF DS to include a direct comparison to a variant without a TXT face. TmAFP was also simulated as a reference. Importantly, we note that our pentapeptide repeat protein simulations, which were prepared using AlphaFold structures for initial coordinates, do not constitute independent structure predictions but rather provide an evaluation of the stability of our AlphaFold structures and indicate whether the relatively subtle differences in the geometric features are preserved as the structures fluctuate in solution.

At 300 K, all of the pentapeptide repeat protein variants remained largely stable for the 100-ns duration of the simulation except for WWWF TT, which showed high CA RMSD values (up to approximately 5 Å) and appeared to be unraveling at the C-terminal end (Figures 3A-D). WWWF TT’s inward residues are especially large and therefore tightly packed, and tryptophans are not found at the central inward positions of interior loops in natural pentapeptide repeat proteins,^54^ making its instability unsurprising. The AlphaFold results had already hinted at this possibility: Only two of WWWF TT’s five predicted structures had passed our filters for analysis; the other three showed the aforementioned topological deviation illustrated in Figure S2C of the Supporting Information, and two of them also had average pLDDT less than 90. In contrast, all five AlphaFold structures for each of LLLL TT, FFFF TT, and FFFF DS had RMSD values less than 1.0 Å with respect to LLLL DS rank 1, and the lowest-ranked structures for those variants had average pLDDT values of 94.4, 96.6, and 95.4, respectively.

**Figure 3:**
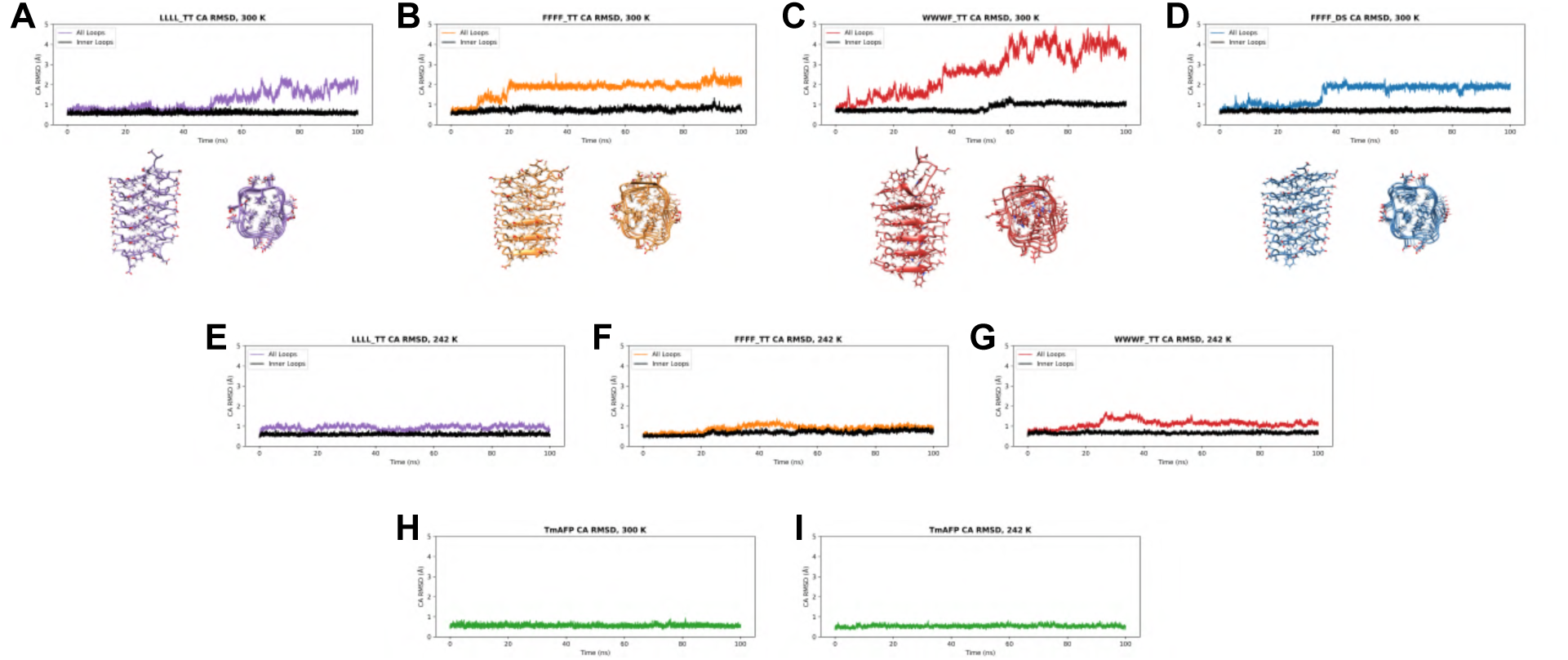
Alpha-carbon RMSD with respect to the initial structure over a 100-ns simulation for (A) LLLL TT at 300 K, (B) FFFF TT at 300 K, (C) WWWF TT at 300 K, (D) FFFF DS at 300 K, (E) LLLL TT at 242 K, (F) FFFF TT at 242 K, (G) WWWF TT at 242 K, (H) TmAFP at 300 K, and (I) TmAFP at 242 K. Below the RMSD plots in each of (A) through (D) are two views of the final structure from the end of the respective simulation at 300 K. In (A) through (G) the black line represents the CA RMSD for only the inner (non-terminal) loops, while the colored line shows the RMSD over all alpha carbons.

We next simulated the three threonine ladder-bearing variants, along with TmAFP, at 242 K, a temperature near the melting point of the TIP4P-Ew water model. ^87,88^ LLLL TT, FFFF TT, and WWWF TT were all reasonably stable at 242 K (Figures 3E-G), and TmAFP showed very little variability at either temperature (Figures 3H and 3I), emphasizing this protein’s significant rigidity, as others have described previously. ^35,89^ We observed for all of the pentapeptide repeat variant simulations at both temperatures that the CA RMSD for only the inner four loops stayed lower than the overall CA RMSD (Figures 3A-G), suggesting that the majority of the structural deviations came from variability in the terminal loops. The relative stability of the inner loops, even in WWWF TT at 300 K, confirms that they remained sufficiently solenoidal over the course of these simulations to permit analysis of the aforementioned geometric features in this subset of our models, and also to investigate whether the threonine-containing variants exhibit IBS-like behavior at low temperature.

While showing differences in the values of some geometric features, the simulations at 300 K preserved the relationships between solenoid variants with different inward residue volumes, as well as the contrast between analogous variants with and without a threonine ladder, that were originally observed in the AlphaFold structures. Compared to these simulations, AlphaFold’s rank 1 structures appeared to underestimate the (i + 8)-(i + 18) CA distance for FFFF DS, FFFF TT, and WWWF TT but not LLLL TT (Figure 4A). However, like in the predicted structures, this distance increased with inward residue volume from LLLL TT to FFFF TT to WWWF TT, and was smaller for FFFF DS than for FFFF TT. The latter outcome is consistent with the threonine-free variants generally falling below the threonine ladder variants in Figure 2A and suggests that the threonines themselves may slightly reduce the concavity of the face they occupy compared to the aspartate/serine combination on the same scaffold. The simulated threonine ladder CB-CA-CB angle distributions also mirrored the predicted dependence on inward residue volume, with WWWF TT nearest to TmAFP’s distribution, LLLL TT farthest from TmAFP, and FFFF TT intermediate (Figure 4B). Interestingly, LLLL TT’s contrast to TmAFP’s IBS in concavity appeared somewhat more drastic in the simulation than in the AlphaFold prediction. FFFF DS exhibited smaller angle values than FFFF TT, again maintaining the trend from AlphaFold.

**Figure 4:**
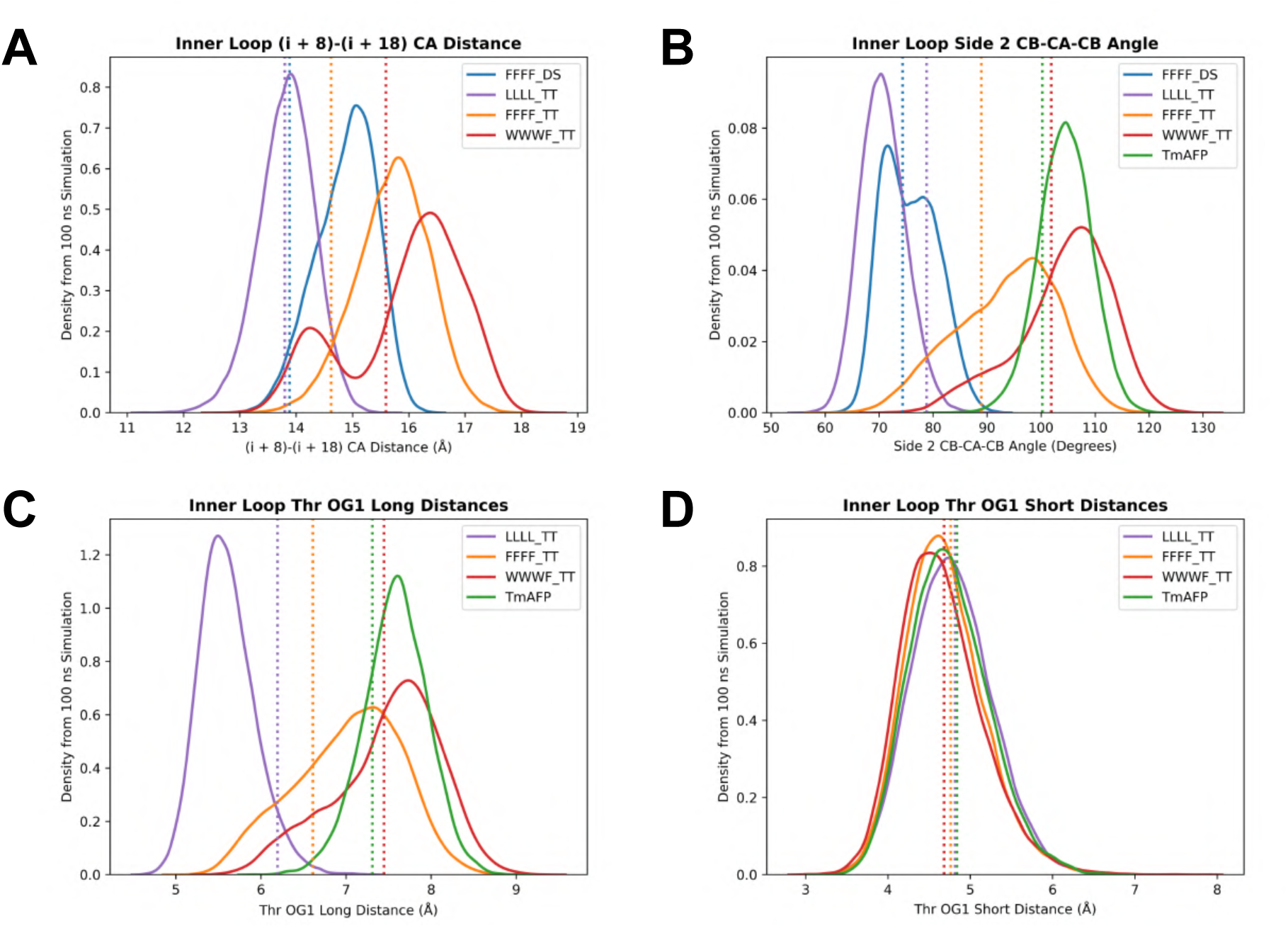
Simulation results at 300 K, including distributions of (A) (i + 8)-(i + 18) CA distances, (B) side 2 CB-CA-CB angles, (C) threonine OG1 long distances, and (D) threonine OG1 short distances. The OG1 distances are filtered based on the threonines’ rotameric states; the unfiltered distributions at 300 K are shown in Figures S3C and S3D of the Supporting Information. Terminal loops are not included in these distributions. In all panels, vertical dashed lines represent the values from the rank 1 AlphaFold structures (or PDB 1EZG chain A for TmAFP).

**Figure 5:**
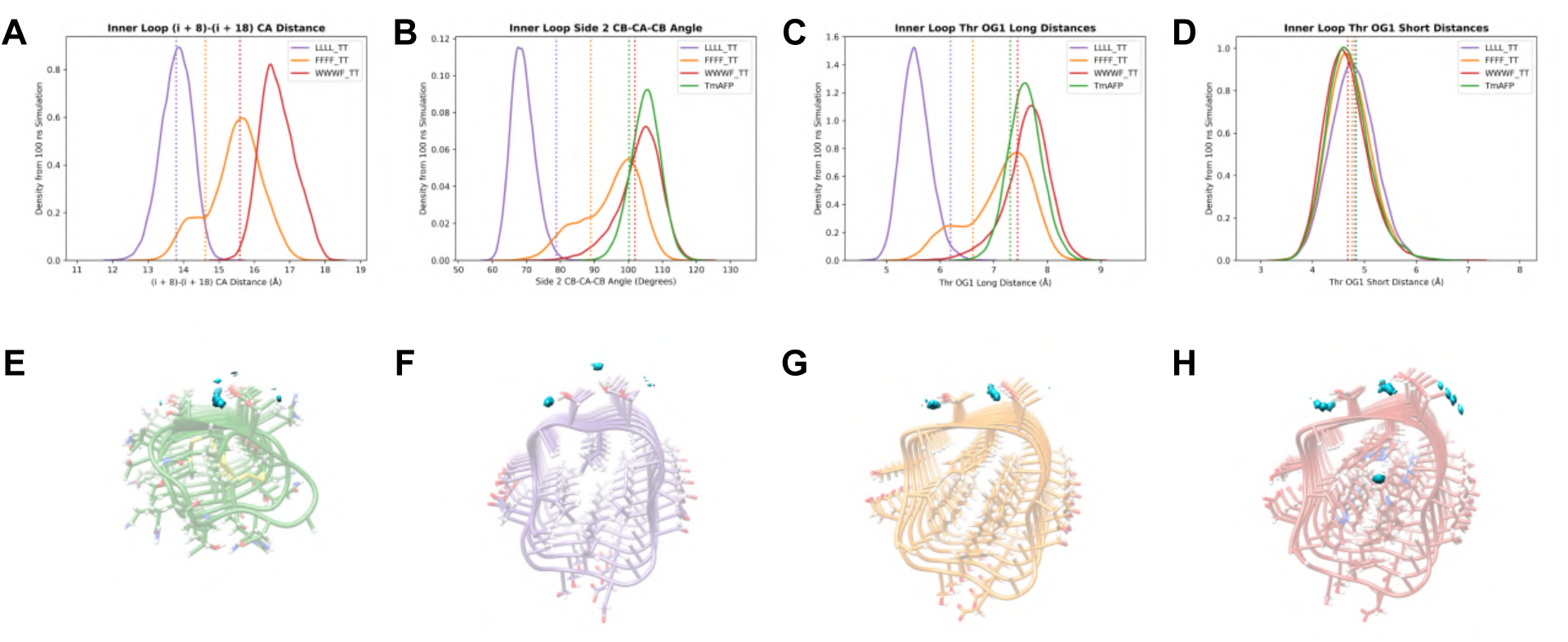
Simulation results at 242 K, including distributions of (A) (i + 8)-(i + 18) CA distances, (B) side 2 CB-CA-CB angles, (C) threonine OG1 long distances, and (D) thre-onine OG1 short distances, again only for inner (non-terminal) loops and with values from the initial structures indicated with vertical dashed lines. The OG1 distances are filtered based on the threonines’ rotameric states; the unfiltered distributions at 242 K are shown in Figures S3E and S3F of the Supporting Information. (E) through (H) show the locations of peaks in water oxygen density around the threonine ladders alongside average structures of (E) TmAFP, (F) LLLL TT, (G) FFFF TT, and (H) WWWF TT with a threshold of 5%. Note that the average coordinates do not necessarily represent realistic individual conformations and some side chains therefore appear distorted.

Analyzing the threonine OG1 distances from the simulations necessitated accounting for variability in the rotameric states of the threonines over time. Consistent with experimental findings^77,78^ as well as previous computational modeling,^36^ in our simulations TmAFP’s threonines predominantly occupied the same rotameric conformation observed in the crystal structure^26^ (Figures S3A and S3B of the Supporting Information). However, the threonines in TmAFP and our pentapeptide repeat variants showed a sufficient degree of rotational variability, particularly at 300 K, to warrant filtering simulation frames based on threonine rotamers to permit a more direct comparison to OG1 distances from our AlphaFold structures and also to focus on the state of the threonine ladders believed to be optimal for ice binding. As it has been suggested that the rotations of the threonine side chains may differ between different protein force fields,^90^ we further reasoned that restricting our distance analysis to the relevant rotameric state provided a simple way to reduce the force field dependence of our conclusions.

Figures 4C and 4D present the threonine OG1 long and short distance distributions at 300 K after this rotamer filtering. WWWF TT and to a somewhat lesser degree FFFF TT substantially overlapped TmAFP’s OG1 long distance distribution, while the simulation reinforced LLLL TT’s contrast (Figure 4C). Like in the predicted structures, the OG1 short distances all displayed very similar distributions in these simulations (Figure 4D). The OG1 distance distributions without rotamer filtering are included in Figure S3 of the Supporting Information; they showed the same trends in peak location with the exception that the filtering was critical to disambiguate a bimodal distribution in TmAFP’s OG1 long distance at 300 K (Figure S3C of the Supporting Information).

Our simulations at 242 K confirmed that the same relationships between the structural features of LLLL TT, FFFF TT, WWWF TT, and TmAFP persisted at a functionally relevant temperature (relative to the water model). At this lower temperature, LLLL TT again exhibited a smaller (i + 8)-(i + 18) CA distance than that of FFFF TT which was still smaller than that of WWWF TT (Figure 5A). WWWF TT followed by FFFF TT continued to be closer to TmAFP than LLLL TT was in both the side 2 CB-CA-CB angle and the threonine OG1 long distance (Figures 5B and 5C). Like at 300 K, the OG1 short distances showed little difference between the proteins (Figure 5D). The OG1 distance distributions without rotamer filtering produced the same relationships with more variability (Figures S3E and S3F of the Supporting Information).

Interestingly, at 242 K FFFF TT’s (i + 8)-(i + 18) CA distance possessed a well-defined lower shoulder, a detail mirrored in the shapes of this variant’s CB-CA-CB angle and threonine OG1 long distance distributions (Figures 5A-C). We supposed that one or more changes between two states of the solenoid’s profile might have impacted the threonine ladder features. To explore this possibility, we assessed the (i + 8)-(i + 18) CA distance and its perpendicular analog, the (i + 3)-(i + 13) CA distance, as a function of time (Figures S4A and S4B of the Supporting Information). Slightly more than 20 ns into the simulation, the former distance increased while the latter decreased: The originally square profile had shifted toward a more rectangular state. At the same point in time, the side 2 (TFT) CB-CA-CB angle increased (Figure S4C of the Supporting Information). Meanwhile the opposing side 4 (DFS) CB-CA-CB angle (Figure S4D of the Supporting Information) showed no distinguishable response to the elongation of the solenoid’s profile in one dimension. These observations suggest that the geometry of a threonine ladder is more tightly coupled to the shape of the solenoid scaffold than is the geometry of a DXS face, corroborating the distinction that appeared in our CB-CA-CB angle data from AlphaFold, with the threonine-free variants lacking the dependence on inward residue volume (Figure 2B).

Our use of explicit water in the simulations permitted analysis of ordered waters near the threonines. Prior studies have established the presence of ordered waters between the threonine rows in several AFPs;^26,31,33,35,36,38,40,41^ we used our low-temperature simulations to determine whether the LLLL TT, FFFF TT, and WWWF TT variants possessed these channel waters. Like TmAFP, FFFF TT and WWWF TT displayed the expected peaks in water oxygen density between the threonines (based on an isosurface threshold identifying positions containing a water oxygen for at least 5% of the production frames), whereas LLLL TT lacked these channel waters (Figures 5E-H). Even at a lower threshold of 1%, these peaks were absent between LLLL TT’s threonines (Figure S5 of the Supporting Information). To visualize the positions of the water oxygen density peaks relative to the solenoids, we had created average structures of the proteins from the simulations. We noticed from the average structures that LLLL TT’s second row of threonines (on the right-hand side in the images) appeared to have the hydroxyl hydrogens pointing away from the beta-solenoid on average, unlike in TmAFP, FFFF TT, and WWWF TT, where these hydrogens pointed down toward the channel waters (Figures 5E-H). To examine this difference in more detail, we checked the CA-CB-OG1-HG1 dihedral angle distributions for the second-row threonines (excluding those in terminal loops) from all four simulations at 242 K. As the average structures had indicated, LLLL TT’s threonine hydroxyl groups displayed a clear divergence in orientation from those of the other proteins (Figure S6 of the Supporting Information). In LLLL TT the hydroxyl groups of the second-row threonines appear to donate hydrogen bonds to water oxygens above the threonine ladder (visible in Figure 5F) rather than within its ranks. LLLL TT’s contrast to the other three solenoids was further emphasized by the different orientation of the backbone carbonyl group of the X residues in the TXT motifs, likely making the carbonyl oxygens unavailable for hydrogen bonding with channel waters (Figure S7 of the Supporting Information).

After running these simulations, we noticed a slight inconsistency in the AlphaFold 2 structures of the LLLL TT variant: In the rank 1 and 2 structures, the backbone carbonyl groups of the X residues in the TXT face point slightly inward, whereas they lie roughly flat on the solenoid’s face in ranks 3 through 5 (Table S3 and Figure S8A of the Supporting Information). Meanwhile AlphaFold 3 consistently gave a more pronounced inward carbonyl orientation (Table S3 of the Supporting Information). To check if our use of the AlphaFold 2 rank 1 structure had impacted the simulation results, we ran an additional simulation at 242 K starting instead from the AlphaFold 2 rank 3 structure. The behavior of the carbonyl groups, as indicated by the average N-CA-C-O dihedral angle (Figure S8B of the Supporting Information), became identical to that in the 242 K simulation of the rank 1 structure as soon as the equilibration restraints were removed, confirming that the outcome was not dependent on the choice of initial structure (Figure S8C of the Supporting Information). The predominant carbonyl orientation in the simulations was closer to that predicted by AlphaFold 3 than to either of the orientations from AlphaFold 2. In contrast, the simulated X residue carbonyl orientation of another solenoid predicted to have a relatively concave TXT face (to be discussed in the following section) remained consistent with the AlphaFold 2 prediction (Figure S8C of the Supporting Information).

### Generation and analysis of AFP-like beta-solenoids with computational methods: A preliminary exploration

Our analysis of hypothetical model beta-solenoids using AlphaFold and MD simulations supported our hypothesis that the structural details of a threonine ladder can depend on the solenoid scaffold. However, these simplified variants of a single scaffold shape served only as a starting point for exploring this topic, which we suspect may be a useful consideration in future endeavors to develop novel beta-solenoid AFPs. We therefore concluded our present study with a walk-through of a protein design strategy aimed at generating initial AFP-like beta-solenoid sequences using deep learning tools, summarized in Figure 6A. This exploration allowed us to examine the impact of distinct solenoid scaffolds on threonine ladder behavior in the context of a computational protein design campaign, while also producing early-stage designs suitable for further analysis and potential refinement into testable AFP candidates.

**Figure 6:**
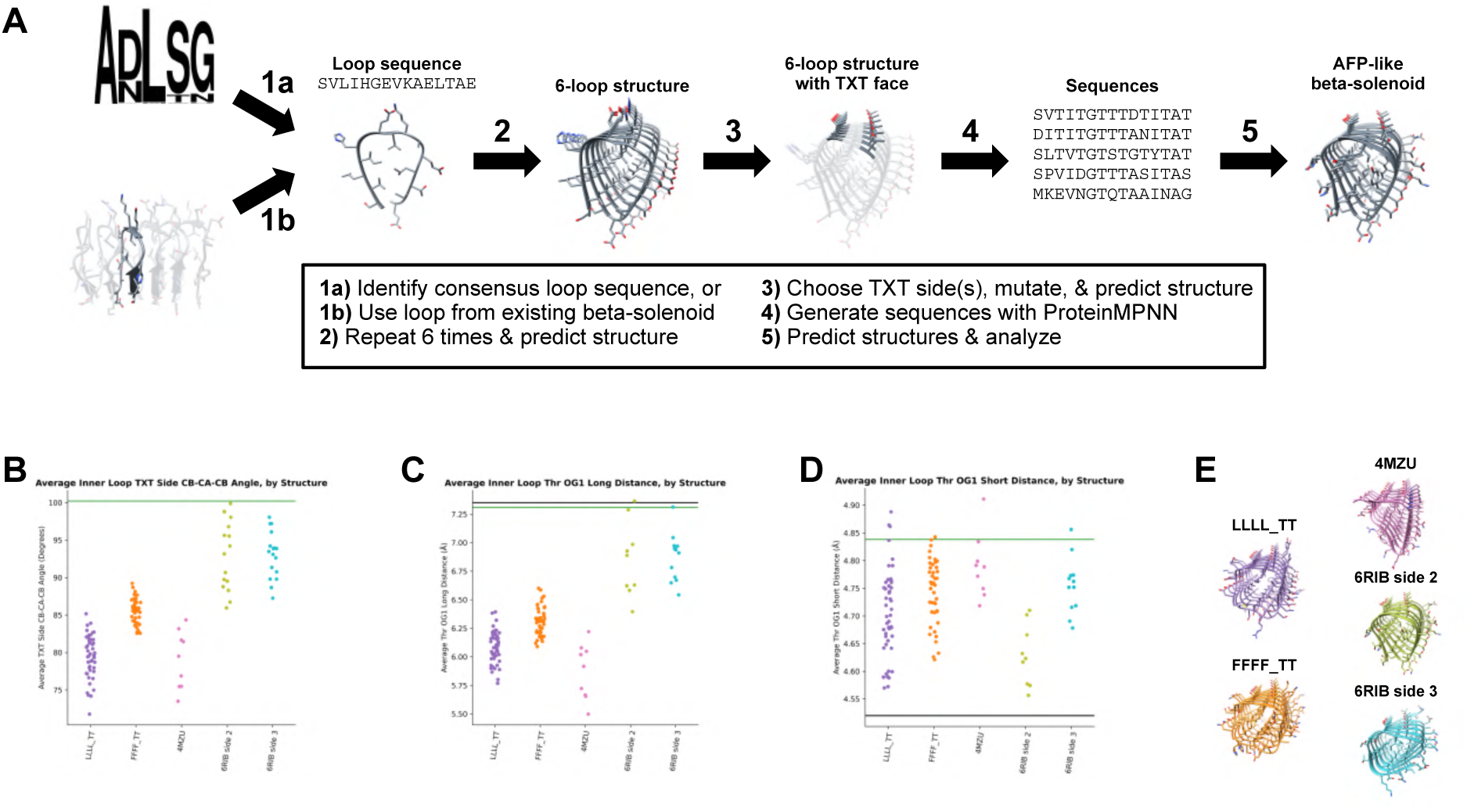
(A) Overview of our strategy to generate beta-solenoid sequence candidates with AFP-like features. (B) Average TXT side CB-CA-CB angles, (C) average threonine OG1 long distances, and (D) average threonine OG1 short distances from non-terminal loops for each predicted structure with pLDDT greater than 90 and CA RMSD less than 2.0 Å with respect to the starting structure that was provided to ProteinMPNN, with (C) and (D) displaying only those that also had all inner loop threonines in the expected rotameric state. The horizontal green lines in (B) through (D) represent values from TmAFP (PDB 1EZG chain A); the horizontal black lines in (C) and (D) represent oxygen spacings from ice: 7.35 Å from the primary prism plane and 4.52 Å from the primary prism or basal plane, respectively.^26^ (E) Rank 1 AlphaFold 2 structures for one selected sequence from each starting point: the candidate with all five predicted structures passing our pLDDT and RMSD filters that had the largest average CB-CA-CB angle. The corresponding sequences are listed in Table S4 of the Supporting Information.

With our initial creation of variants from the pentapeptide repeat consensus ADLSG, we already established that predicted beta-solenoid structures—potentially useful for subsequent design steps—can be derived from consensus sequences. We supplemented our approach by exploring an alternative means of constructing possible AFP scaffolds based on single loops of individual beta-solenoids from nature. Assisted by Kajava and Steven’s thorough inventory of beta-solenoid proteins,^91^ we browsed the Protein Data Bank^92^ and selected 4OJ5,^93^ 4MZU,^94^ and 6RIB^95^ as starting points. These structures, with distinct profile shapes and varying degrees of regularity, represent a putative bacteriophage tailspike protein, a bacterial biosynthetic enzyme, and a bacterial cytoskeleton component, respectively. From each structure we identified the sequence of one arbitrarily chosen loop of the solenoid; a second, different loop was taken from 4MZU for greater variety. Similar to our original strategy of repeating ADLSG to form a regular six-loop solenoid, we repeated each loop sequence from the PDB structures six times to produce initial solenoid sequences.

We predicted five structures for each initial solenoid with AlphaFold 2. The solenoid derived from 4OJ5 had a large and somewhat variable profile, with long faces that could be suitable for hosting more than two rows of threonines (Figure S9 of the Supporting Information); while less relevant to our current investigation centered on the two-row threonine IBS, scaffolds like this one may yet hold promise for future design of AFPs/IBPs with larger ice-binding faces. This solenoid was not subjected to further engineering in the present study. One of the two sequences from 4MZU was also dropped, having failed to produce six-loop beta-solenoid structures: The top-ranked prediction was a three-loop beta-solenoid with an average pLDDT of 58.0 and the four lower-ranking structures contained alpha helices. Intriguingly, the 6RIB loop gave five structures with at least relatively high average pLDDT (ranging from 80.4 to 95.6) but only three were right-handed beta-solenoids; the rank 3 structure was a single alpha helix and the rank 5 structure was a left-handed beta-solenoid. This example bears some resemblance to a case described by Pratt et al. ^84^

As the other 4MZU loop and the 6RIB loop had both yielded confident six-loop solenoid structures (albeit with some variability), we chose to move forward with generating designs based on these solenoid shapes. We used their highest-ranked predicted structures to visually select faces with two rows of outward-facing residues to bear the threonine ladder as a potential IBS: We proceeded with one face for the solenoid from 4MZU and two different faces for the solenoid from 6RIB (labeled side 2 and side 3). We mutated the sequences accordingly and predicted structures again. At this point we had produced three perfectly repetitive, highly regular predicted beta-solenoids with threonine ladders, analogous to the LLLL TT, FFFF TT, etc. variants we had developed from the consensus sequence. These presented desirable structures but not necessarily desirable sequences.

To diversify these repetitive beta-solenoids and develop potentially more promising sequences, we next used the deep learning model ProteinMPNN to generate sequence candidates conditioned on the rank 1 AlphaFold structures of the PDB-derived solenoids (from 4MZU and 6RIB side 2 and side 3) and the consensus-based LLLL TT and FFFF TT solenoids. Initial tests with LLLL TT suggested that a high sampling temperature is necessary to achieve substantial variability in generated beta-solenoid sequences (Figure S10 of the Supporting Information). Thus with a sampling temperature of 0.8, we generated 10 sequences for each of the five structures, keeping the putative IBS threonines fixed and prohibiting cysteines. AlphaFold 2 structures were predicted for all of the sequences. While a small number of sequences per structure was sufficient for the purpose of this illustration, the computational efficiency of ProteinMPNN and AlphaFold makes this approach readily applicable to larger batches of designs.

Even at this high sampling temperature, ProteinMPNN produced many sequences confidently predicted to form beta-solenoids. Out of 250 structures (five predicted structures for each of 10 sequences for each of five starting structures), 132 had average pLDDT greater than 90 and showed CA RMSD less than 2.0 Å relative to their respective starting structure that was provided to ProteinMPNN. Those 132 structures represented 30 different sequences, 21 of which had all five predicted structures passing the pLDDT and RMSD cutoffs (Table 1). Only the 6RIB side 2 batch yielded entirely distinct structures with moderate confidence (Figure S11 of the Supporting Information).

**Table 1:**
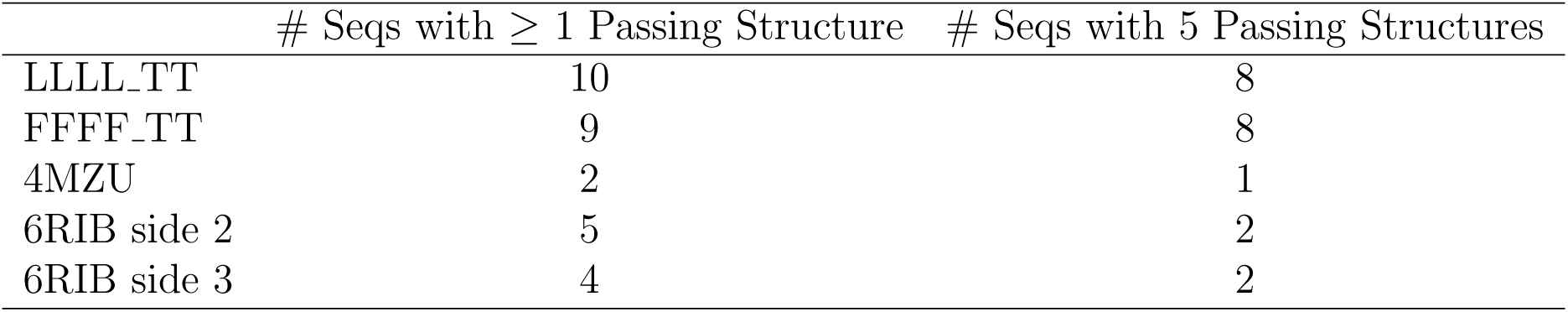
AlphaFold 2 Results for ProteinMPNN Sequences.

| | # Seqs with $\geq 1$ Passing Structure | # Seqs with 5 Passing Structures |
| --- | --- | --- |
| LLLL_TT | 10 | 8 |
| FFFF_TT | 9 | 8 |
| 4MZU | 2 | 1 |
| 6RIB side 2 | 5 | 2 |
| 6RIB side 3 | 4 | 2 |

We compared the CB-CA-CB angle of the TXT face for all passing structures and the threonine OG1 distances for all structures with inner loop threonines in the proper rotameric state (Figures 6B-D). Consistent with our expectations based on the model solenoids, in these predictions the scaffold choice appeared capable of influencing local structural details of the threonine ladder, confirming that scaffold-dependent differences in threonine behavior may be encountered in a beta-solenoid design process. Like in our original AlphaFold results and MD simulations, the CB-CA-CB angle and OG1 long distance produced similar patterns, with each showing the same relationships between the five groups of designs; using both of these features to compare designs may therefore be somewhat redundant. Furthermore, the OG1 long distance again varied over a larger range than did the short distance. Between the LLLL TT and FFFF TT starting structures, FFFF TT yielded designs with CB-CA-CB angle and OG1 long distance values closer to TmAFP’s, implying that ProteinMPNN had at least in some instances preserved subtle differences between the initial structures. The designs based on 4MZU exhibited small angles and OG1 long distances comparable to those from LLLL TT, indicating that these two scaffold sets possessed the most concave TXT faces. The sequences produced from 6RIB (both side 2 and side 3) came nearest to TmAFP in both angle and long distance values and thus closest to the aforementioned longer ice lattice spacings, suggesting that this scaffold shape may offer multiple options for the location of a possible IBS with similar effects on threonine ladder geometry.

Having previously observed in our simulations the instability of a variant (WWWF TT) that had variable AlphaFold results, we reasoned that the ProteinMPNN sequences with the most consistent and confident predicted structures may have the greatest likelihood of stability. For each group of designs, we therefore consider the sequences with all five AlphaFold structures passing our pLDDT and RMSD filters to be the most promising for further analysis. Figure 6E displays the top-ranked AlphaFold structure for the sequence from each batch that met those criteria and had the largest CB-CA-CB angle (averaged over all five structures); the sequences are provided in Table S4 of the Supporting Information. The CB-CA-CB angle appeared particularly useful for comparing designs because it had correlated with the scaffold-sensitive OG1 long distance’s similarity to ice lattice spacings while being less susceptible to variability in threonine rotamers.

As three of our resulting sequence groups (the 4MZU, 6RIB side 2, and 6RIB side 3 batches), were not derived from pentapeptide repeat scaffolds and therefore differed substantially from the solenoids we had analyzed so far, we ran low-temperature simulations of the selected designs from these groups to obtain additional data on these scaffolds. The simulations were consistent with AlphaFold’s suggestion that the 6RIB-based designs came nearer to TmAFP in CB-CA-CB angle and OG1 long distance values than did the 4MZU-based design, which was closer to (but still above) the distributions from the original LLLL TT model solenoid (Figures S12A and S12B of the Supporting Information) and did not exhibit inward-facing carbonyl groups (Figure S8C of the Supporting Information). Meanwhile the OG1 short distances continued to show less sensitivity (Figure S12C of the Supporting Information). All three proteins exhibited channel water density peaks within their threonine ladders (Figure S12D-F of the Supporting Information), confirming that our design strategy can yield predicted structures that reproduce this hallmark of natural ice-binding threonine surfaces in simulations. These preliminary designs appear suitable for further investigation and refinement, which are beyond the scope of our current study centered on analysis of threonine ladder differences. In this way, we have illustrated the relevance of our findings from the pentapeptide repeat variant models in a computational protein design context and have taken early steps toward developing novel, cysteine-free beta-solenoid designs that may eventually merit experimental assessment of ice growth modulation.

## Discussion

The threonine ladders found on a subset of AFPs are perhaps nature’s most elegant solution to what has been described as the greatest recognition challenge in biology — the differentiation between water’s solid and liquid phases.^96^ Two rows of threonine residues constitute a streamlined IBS that is sufficient for ice recrystallization inhibition at submicromolar concentrations^97^ as well as high thermal hysteresis activity,^28,29^ making this type of IBS very promising for use in designing novel AFPs. By demonstrating that the local structure of a threonine ladder can be significantly influenced by the beta-solenoid scaffold, we attempted to augment the growing body of knowledge of IBS function that will inform design efforts in this area.

Our findings not only reinforce past comparisons but also have both retrospective and prospective implications for AFP design. Scaffold-dependent differences in the concavity of the solenoid faces might be a factor contributing to disparities in the thermal hysteresis activity^62^ and computationally studied water ordering^40,61^ between insect AFPs and bacterial INP segments. Additionally, our comparisons indicate one possible reason for the low thermal hysteresis activity of the pentapeptide repeat protein that was previously transformed into a new AFP by Yu.^57^ The cyanobacterial pentapeptide repeat protein used (PDB 2J8K) had the LLLL pattern of inward-facing hydrophobes throughout all of its loops, with the exception of a single phenylalanine near the N-terminal cap.^98^ Based on the analysis of our LLLL TT model variant, this solenoid’s installed TXT face may have been overly concave, with small OG1 long distances, and possibly unable to accommodate channel waters as well as a natural AFP would. Intriguingly, the ice shaping that was nevertheless observed for Yu’s mutated protein hints that even an apparently suboptimal threonine ladder face may not entirely preclude ice binding.

Our results suggest that the activity of this novel AFP might potentially be improved by mutating some of its inward-facing leucines to phenylalanines, or by starting from a pentapeptide repeat protein with higher average inward residue volume in the first place, such as MfpA from *Mycobacterium tuberculosis* which has more internal phenylalanines.^55^ Either option may confer a more AFP-like threonine ladder IBS. It remains unclear whether such dramatic differences as the absence of channel waters may appear frequently in future AFP engineering efforts or whether pentapeptide repeat proteins with internal leucines represent an uncommon extreme case. Nevertheless our observations were sufficient to illustrate that attention to a solenoid scaffold’s impact on threonine hydroxyl spacings, X residue carbonyl behavior, and channel waters may be informative in such efforts.

Looking forward, we expect state-of-the-art methods for protein design to open up many exciting possibilities for IBP design that go beyond mutating natural proteins. For instance, de Haas et al. recently reported compelling results from the application of *de novo* design tools to the creation of alpha-helical IBPs.^99,100^ As the beta-solenoid fold is capable of producing relatively flat, parallel beta-sheet faces with a high degree of regularity, methods for designing beta-solenoids (e.g., Refs. 56,101–103) may be central to the further development of novel ice binders. In the process of considering how different solenoid scaffolds can affect threonine ladder geometry in the context of a computational protein design campaign, we illustrated one potential way to generate new AFP-like beta-solenoid sequences. Our strategy has some conceptual similarities to the beta-solenoid design approaches of Saragovi et al. ^102^ and Pretorius et al. ^103^ but with key differences including: (1) We take starting points from natural beta-solenoid sequences for initial structure building, and (2) we particularly focus on the inclusion of the threonine ladder motif. As we showed in Figure 6 and Figures S11 and S12 of the Supporting Information, this strategy can produce sequences confidently predicted by AlphaFold to fold into regular solenoids with TXT faces that exhibit certain IBS-like properties in simulations; ultimately experiments will be necessary to validate our approach. Furthermore, the applications of methods for beta-solenoid design reach beyond ice modulation into the area of nanomaterials, such as nanowires,^104,105^ cages,^56,106^ solid coatings,^107^ semiconductor templates,^102^ and protein-based porous or catalytic frameworks.^56,108–111^

Additional steps may be necessary to complete our designs. Without capping structures or terminal irregularities, beta-solenoid AFPs can assemble into amyloid fibrils.^104,105^ While forming an extended IBS through self-assembly may be desirable for ice nucleation activity,^112–115^ designs that do not assemble will be more similar to monomeric natural AFPs. Thus we would consider modifying our ProteinMPNN designs by attaching capping groups based on natural solenoids,^56^ diffusing caps with RFdiffusion,^48^ or simply mutating select residues in both terminal loops to like-charged amino acids.

We chose to focus our current investigation on features related to the flatness or concavity of the beta-solenoid face hosting a threonine ladder, but as we acknowledged, this is only one of many factors that may influence an AFP’s ice-binding capacity and activity. We observed in our simulations for instance that TmAFP displayed exceptional backbone rigidity (Figures 3H and 3I), as has been noted previously,^35,89^ which likely contributes to its function; other solenoid scaffolds may be more flexible. Qualitatively, we also noticed subtle differences in our solenoids’ degree of twist (visible in the structures in Figures S12D-F of the Supporting Information). Twisting may reduce a threonine ladder’s complementarity to the ice surface. Aside from the IBS itself, the impact of an AFP’s shape and non-IBS surface chemistry on its activity is only beginning to be understood. ^116^ Our strategy for generating beta-solenoid sequences offers a path to new comparisons that may help to address this topic. Indeed, the expanding toolkit for protein design presents new opportunities to isolate the effects of specific features, as de Haas et al. illustrated by precisely constraining the twist in their alpha-helical IBPs.^99^ Also noteworthy, our hypothetical WWWF TT model with threonine ladder geometry very similar to TmAFP but with evident stability issues highlights that considerations of IBS optimization may perhaps be a secondary priority to that of achieving correctly folded beta-solenoids, as the threonine behavior is of little consequence if the protein will not fold as intended.

It is essential to note the limitations of both of the computational methods we employed (structure predictions and MD simulations). For example, the residue flips observed variably in a subset of our AlphaFold structures, as illustrated in Figure S2C of the Supporting Information, suggest that AlphaFold may easily make local topological errors in beta-solenoid structures even while maintaining high confidence. This type of topological deviation was previously noted when an AlphaFold structure of an INP^60^ was compared to an earlier homology-based model;^59^ Pal et al. recently demonstrated that the difference between the two structures altered the INP’s ice-nucleating ability in simulations, with the AlphaFold structure appearing more favorable. ^61^ Additionally, the inconsistent handedness we observed in the lowest-ranked structure for the initial six-loop sequence from 6RIB suggests that AlphaFold might in some cases struggle to distinguish right-handed from left-handed beta-solenoids. Both cases are represented among natural hyperactive AFPs,^26,69^ thus one handedness may not be strongly preferable in AFP design, but predicting handedness incorrectly may still affect the success of the design process.

Beyond these structural details that may be predicted imperfectly, it has been noted that AlphaFold (particularly AlphaFold 2) can frequently predict confident but unrealistic beta-solenoids for meaningless random sequence repeats, ^84^ prompting us to examine how this capacity to hallucinate improbable solenoids may impact our results and conclusions. Our initial 40 pentapeptide repeat protein variants were derived from a consensus sequence for a family of natural proteins with multiple representatives in the Protein Data Bank (e.g., PDB 2BM4^55^ and 2J8K^98^), thus the predominance of quadrilateral, right-handed beta-solenoids predicted for our sequences is expected and we know this fold can exist for sequences relatively similar to ours. In addition, design efforts have already established the resilience of this fold to redesign beyond the natural sequence space.^56^ That said, the model solenoid sequences we constructed contained inward-facing residues valine or tryptophan that are not well represented at those positions in natural pentapeptide repeat proteins,^54,56^ making it highly plausible that some of the predicted solenoids are fictitious. We observed the most structural variability at the high and low extremes of the inward residue volume range (Figure S2A of the Supporting Information). This outcome is consistent with the inward residue options of intermediate size (leucine and phenylalanine) being those commonly found in natural sequences in the pentapeptide repeat family, ^54^ hinting that at least in aggregate AlphaFold favored the more evolutionarily realistic sequences, even without using multiple sequence alignments. Ultimately, however, our conclusions are not dependent on the model beta-solenoids’ realism; our goal in creating the pentapeptide repeat variants was to analyze potential distinctions between threonine ladders on scaffolds that differed in a specific, controllable way. Analysis even of the particularly unrealistic WWWF TT model variant had value in illustrating that a TXT face with minimal concavity, essentially bulging, enforces ice-like hydroxyl group spacings like those of TmAFP.

On the other hand, AlphaFold’s propensity to predict beta-solenoids has pronounced implications for the design of new AFPs. On the positive side, this propensity makes it easy to generate predicted solenoid designs *in silico*, as we did per Figure 6A. Hallucination of false structures is an acceptable and even useful component of the design process at step 2, where a solenoid structure is created for later use in conditional sequence generation, analogous for instance to the backbone generation element of design campaigns featuring RFdiffusion followed by ProteinMPNN.^48^ It is at step 5 in our illustrated workflow where unrealistic solenoid predictions become problematic: Screening designed sequences for an intended solenoid fold with AlphaFold may be prone to false positives, necessitating additional evaluation of the designs with different computational tools and eventually with experimental assessment of multiple candidates. Despite the possibility of false positives, AlphaFold 2 remains a valuable first step for screening large batches of sequence designs as it is a computationally efficient means of identifying candidates worthy of further attention and, as we have shown, can hint at both potential instability and subtle differences in threonine ladder geometry.

As protein structure prediction methods continue to emerge and improve, ^49,50,52^ future studies will likely explore how they compare to AlphaFold 2 in their capacity to meaningfully predict beta-solenoid structures. Pratt et al. already noted AlphaFold 3’s improvement over AlphaFold 2 in this regard. ^84^ AlphaFold 3 (with default settings) consistently predicted beta-solenoids with fairly high to very high confidence for our selected ProteinMPNN sequences (those in Figure 6E and in Table S4 of the Supporting Information), with little difference from the starting backbone for all except that from 4MZU, which flipped handedness (Figure S14 of the Supporting Information). We note that even with support from AlphaFold 3, false positives are a concern. For the WWWF TT variant that was unstable in our simulation at 300 K, AlphaFold 3 still predicted the quadrilateral beta-solenoid fold, with slightly lower confidence (average CA pLDDT ranging from 81.9 to 83.3) compared to AlphaFold 2 (average pLDDT ranging from 83.1 to 94.0) but less structural variability (no flipped alanines).

MD simulations can also be employed in subsequent filtering and analysis of beta-solenoid AFP designs that appear promising based on structure predictions. However, simulations come with their own set of limitations. Our 100-ns simulations may not have been long enough to reveal instability in all cases, allowing some structures to appear stable when they may actually be only metastable. Inaccuracies in common protein force fields and water models can further limit the predictive ability of simulations, and different force fields and water models can yield entirely different conclusions about structural and dynamical characteristics.^90,117–122^ We controlled for this impact on threonine rotamer variability but not in other contexts. For instance, while our low-temperature simulation of TmAFP showed its second-row threonine hydroxyl hydrogens pointing down toward the channel waters (Figures 5E and S6 of the Supporting Information), in a recent study Zhang et al. opted for different protein and water parameters and displayed simulation snapshots with the hydroxyl hydrogens pointing up, similar to our LLLL TT case (Figure 5F), but with the channel waters still present.^41^ More accurate models for simulating proteins in water will be necessary to resolve this ambiguity. Nevertheless, our simulations provided a physics-based corroboration of the trends that emerged in our structure predictions and indicated how ordered waters may fit into these trends.

## Conclusions

In the present work we set out to provide new details to inform the future design of beta-solenoid AFPs. We conclude that the geometry of a threonine ladder is likely sensitive to the beta-solenoid scaffold upon which it is arrayed, and that relatively subtle differences between scaffolds can be sufficient to impact the threonines’ ability to match the ice lattice and even to alter the structure of local hydration water. Our exploration of specific AFP design considerations fits into a broader landscape of recent advances^84,101–103^ underscoring that beta-solenoids may be a challenging yet valuable special case for computational protein design. We hope our findings, combined with continuing progress in structure prediction tools, protein design methods, and simulation potentials, will support efforts to develop a new generation of highly active AFPs suitable for various practical applications requiring the modulation of ice growth.

## Supporting Information Available

Additional figures showing structure prediction and simulation results and observations from the beta-solenoid design demonstration, as well as tables containing all model solenoid loop sequences, LLLL TT carbonyl orientation data, and selected ProteinMPNN sequences, and representative inputs for our Amber simulations.

## Acknowledgement

We thank Konrad Meister, Ilja Voets, and Renko de Vries for insightful discussions of ice-binding proteins and beta-solenoids, as well as all members of the Paesani Lab for continuous support and feedback. This research was supported by the Air Force Office of Scientific Research under award no. FA9550-20-1-0351. Computational resources were provided by the Department of Defense High Performance Computing Modernization Program (HPCMP). C.N.C. was partially supported by the San Diego Fellowship.

---

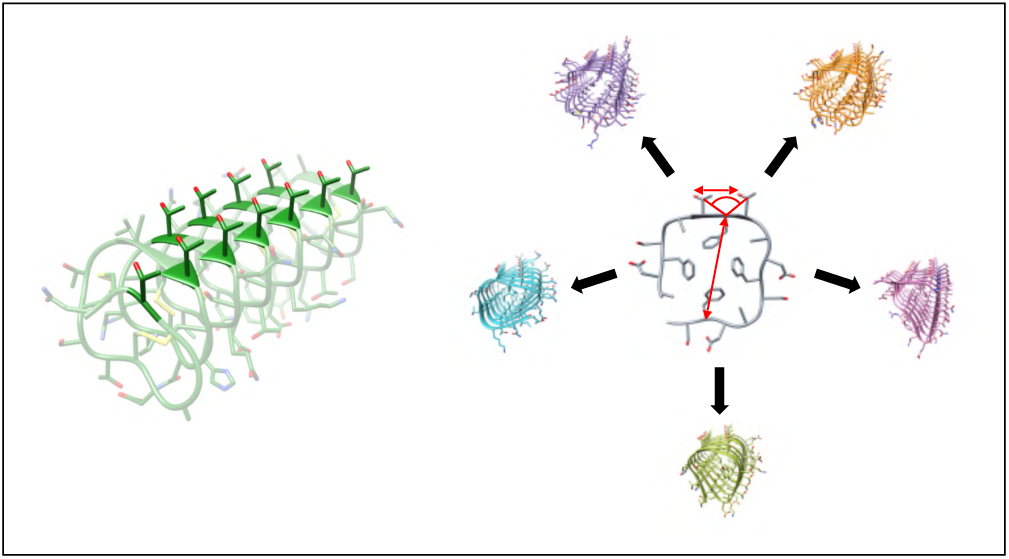
TOC Graphic

