## Supporting Information for "Computational analysis of threonine ladders on distinct beta-solenoid scaffolds, with implications for the design of novel antifreeze proteins"

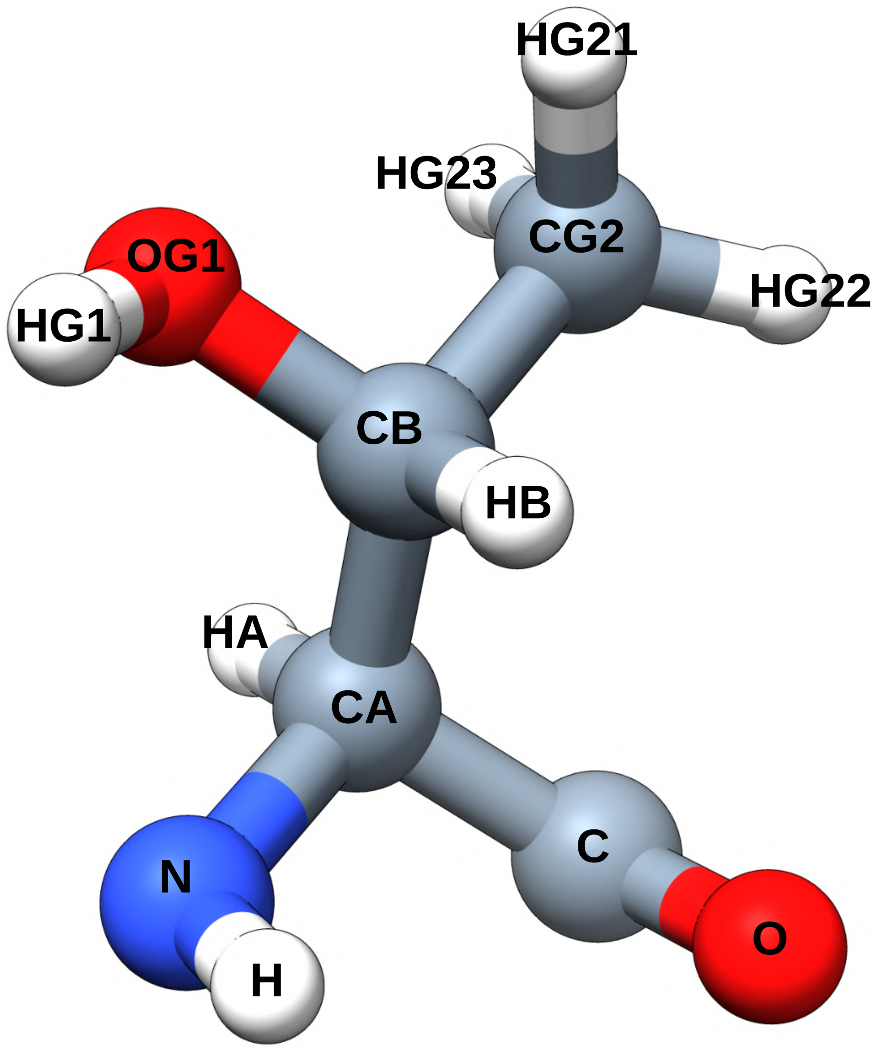


Figure S1: Atom names, as used throughout our analysis, for a threonine residue.

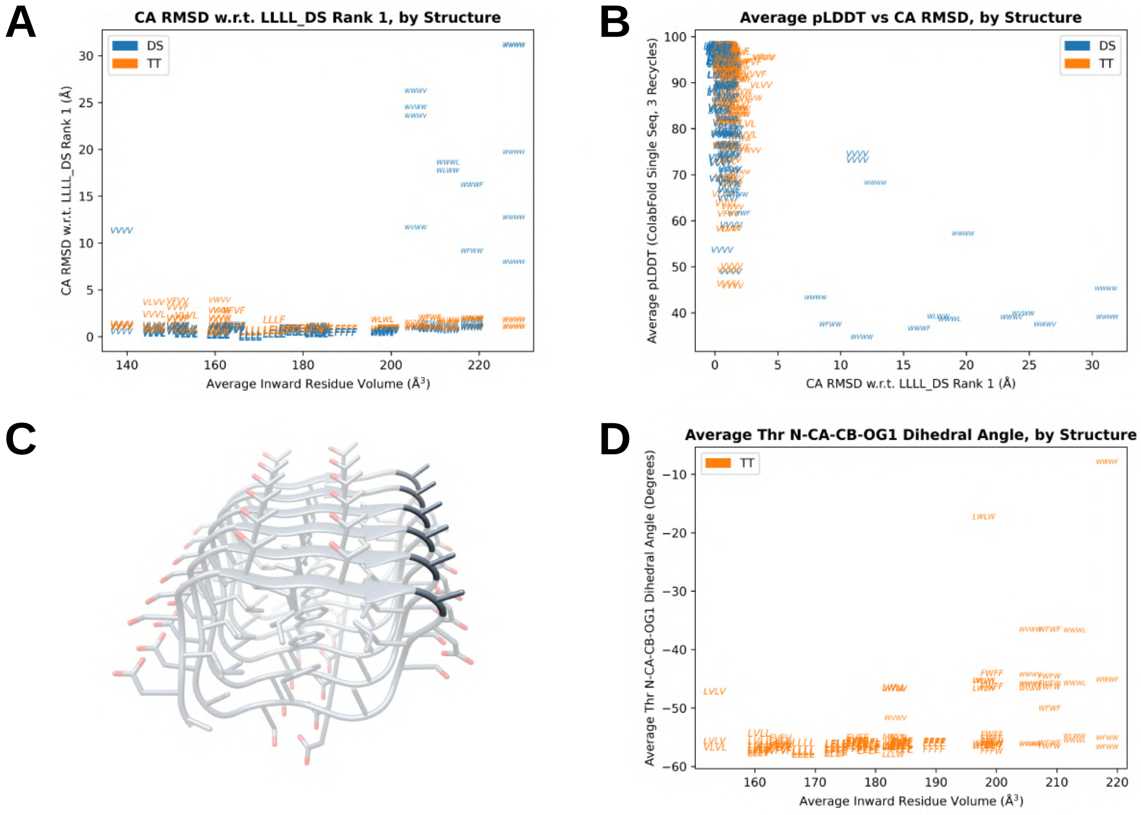


Figure S2: (A) Alpha-carbon RMSD of all pentapeptide repeat variant AlphaFold structures (with respect to the top-ranked structure of LLLL_DS) versus the average inward residue volume. (B) Average pLDDT versus CA RMSD (again with respect to LLLL_DS rank 1) for all pentapeptide repeat variant AlphaFold structures. (C) Example of a predicted structure in which a corner alanine (darkened in the image) is flipped outward in each loop, deviating from the typical square profile. (D) Average N-CA-CB-OG1 dihedral angle over all loops for each predicted TXT variant structure with average pLDDT greater than 90 and CA RMSD to LLLL_DS rank 1 less than 2.0 Å and without topological deviations.

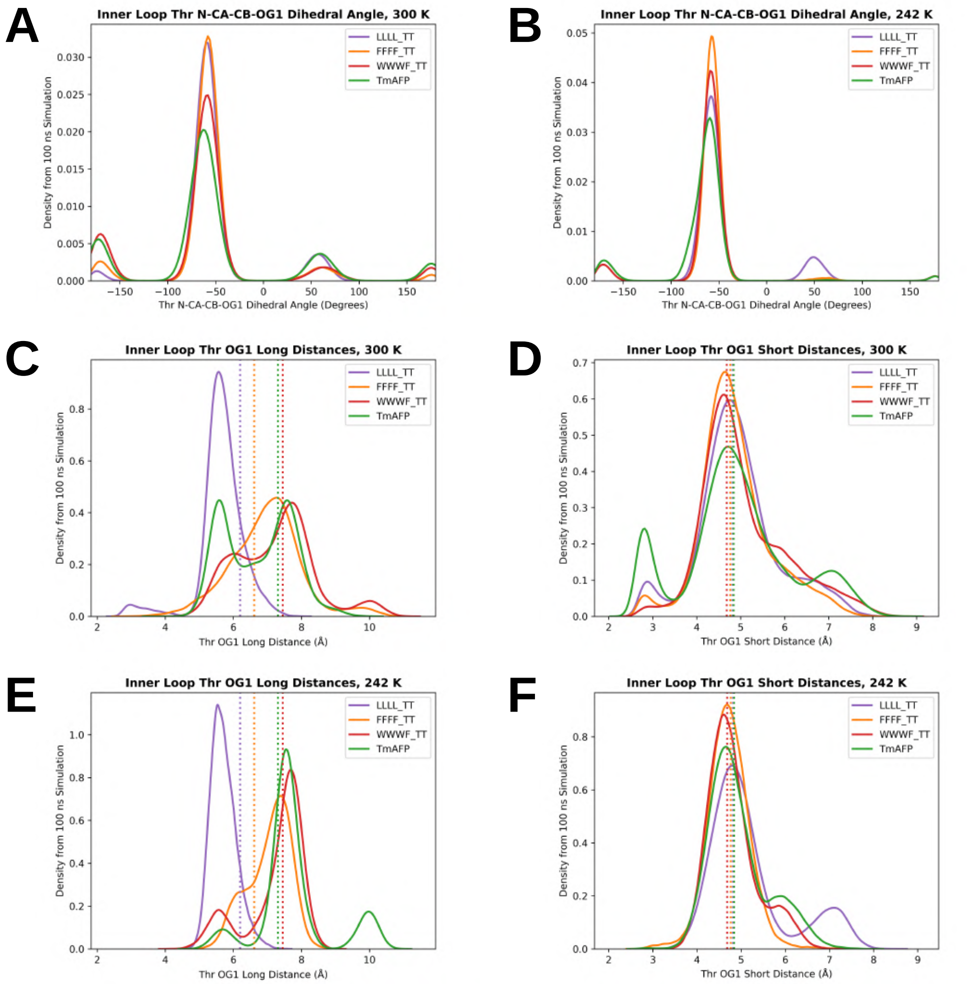


Figure S3: Threonine N-CA-CB-OG1 dihedral angle distributions from simulations at (A) 300 K and (B) 242 K, including only inner loops. Unfiltered inner loop threonine OG1 (C) long and (D) short distance distributions at 300 K, and (E) long and (F) short distance distributions at 242 K. In (C) through (F), vertical dashed lines represent the values from the rank 1 AlphaFold structures (or PDB 1EZG chain A for TmAFP).

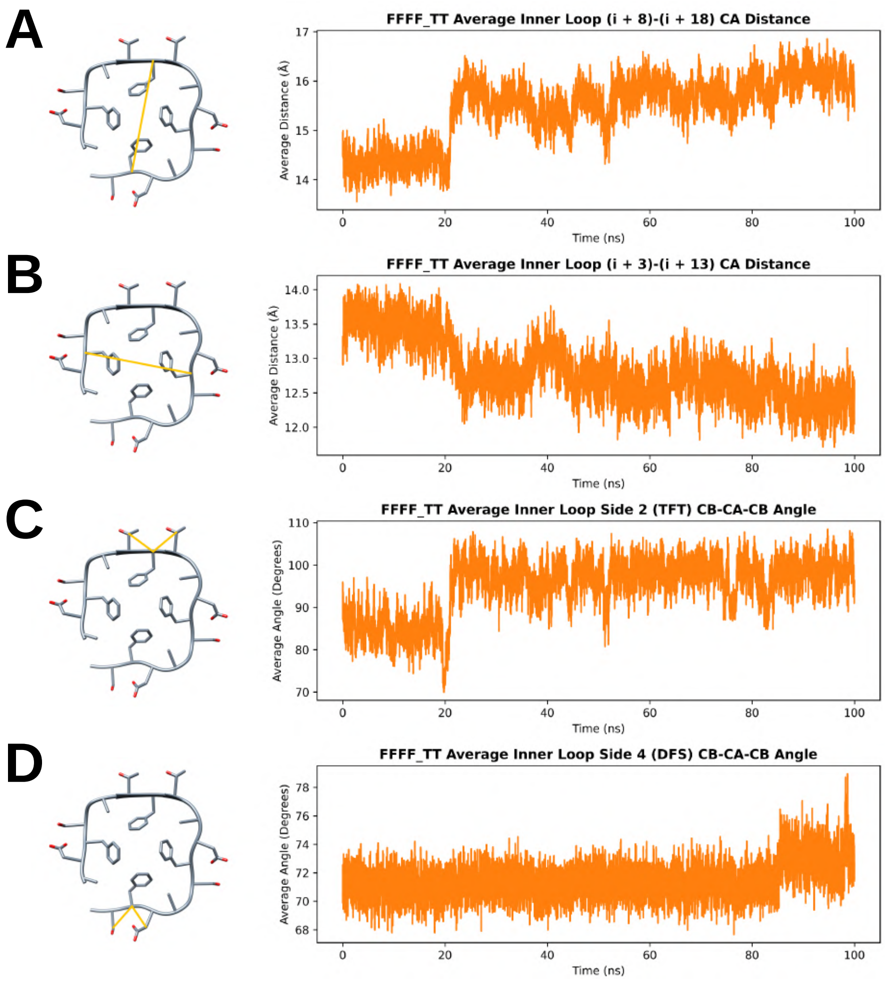


Figure S4: Average inner loop (A) (i + 8)-(i + 18) CA distance, (B) (i + 3)-(i + 13) CA distance, (C) side 2 TFT CB-CA-CB angle, and (D) side 4 DFS CB-CA-CB angle versus time from the simulation of FFFF_TT at 242 K. Each of these geometric features is illustrated to the left of the corresponding plot.

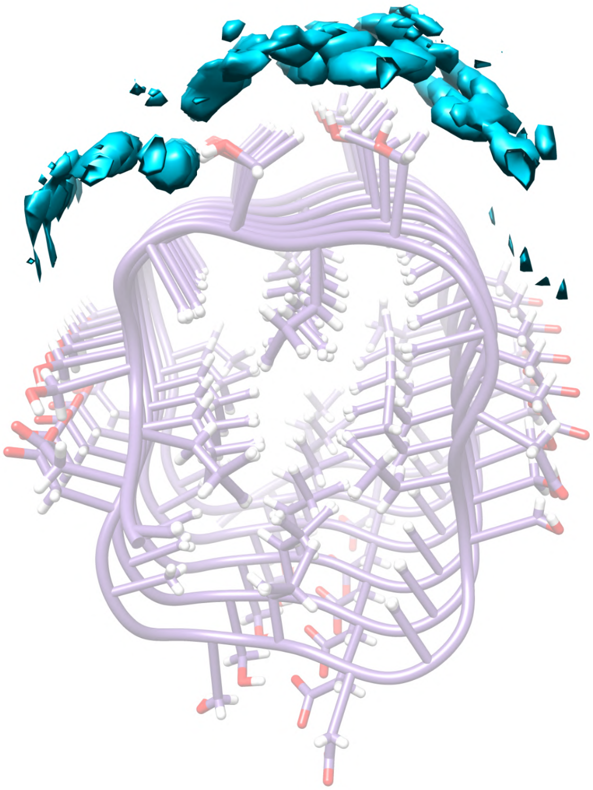

Figure S5: Water oxygen density distribution around LLLL_TT's threonine ladder, shown with the average structure, from the simulation at 242 K, with a threshold of 1%.

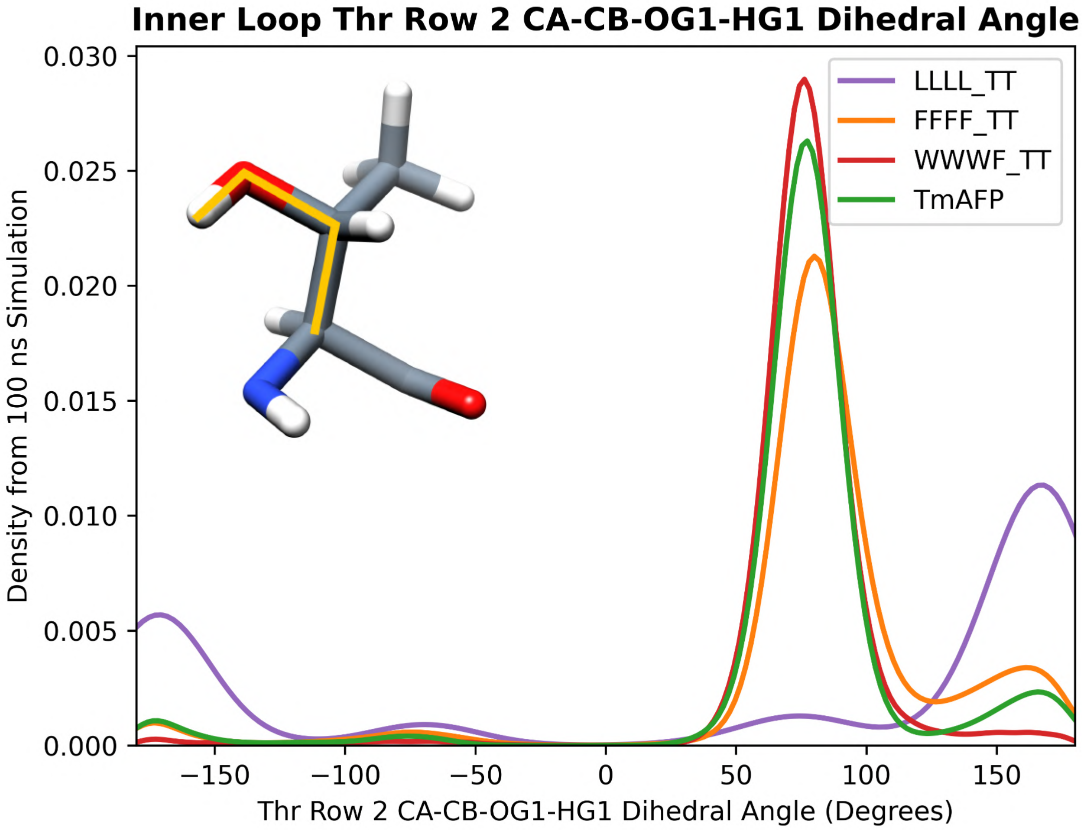


Figure S6: Distributions of the CA-CB-OG1-HG1 dihedral angles (inset) from second-row threonines in inner loops at 242 K.

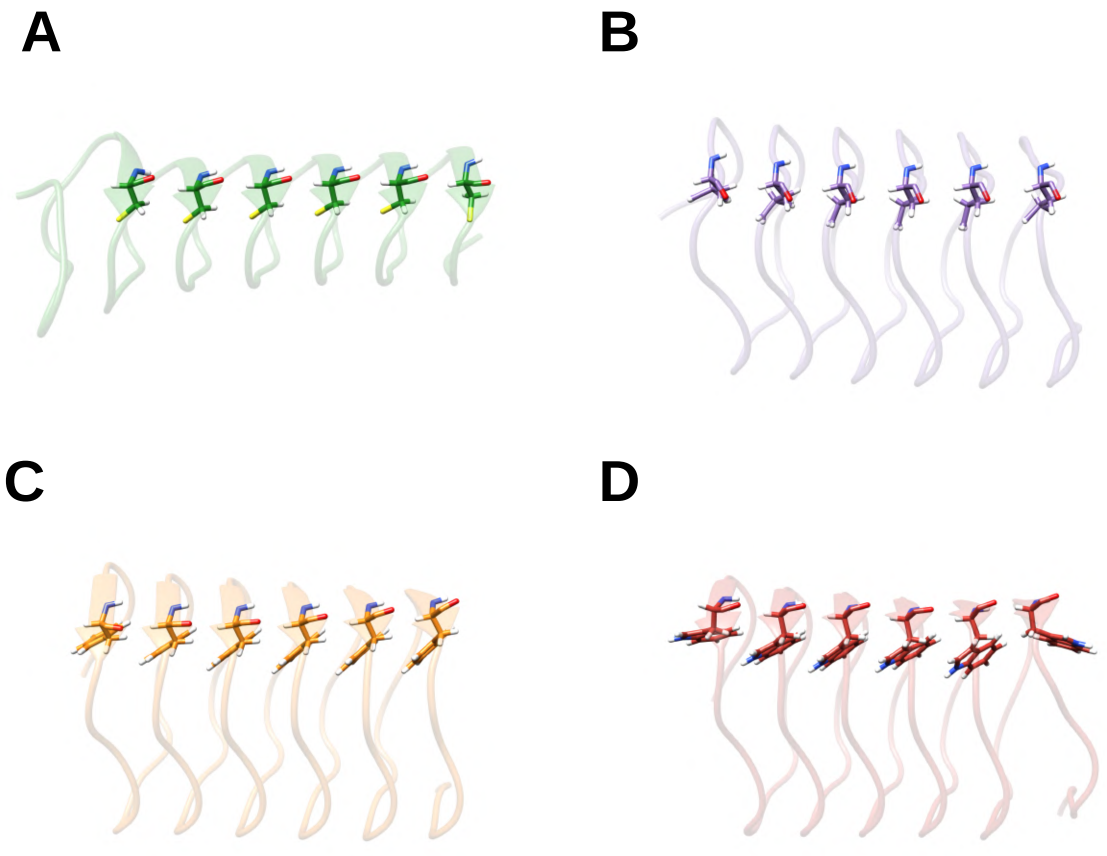


Figure S7: Average structures of (A) TmAFP, (B) LLLL_TT, (C) FFFF_TT, and (D) WWWF_TT from the simulations at 242 K, with average atom positions shown only for the inward-facing X residue in each of the TXT motifs (or AXT for the loops near TmAFP's termini).

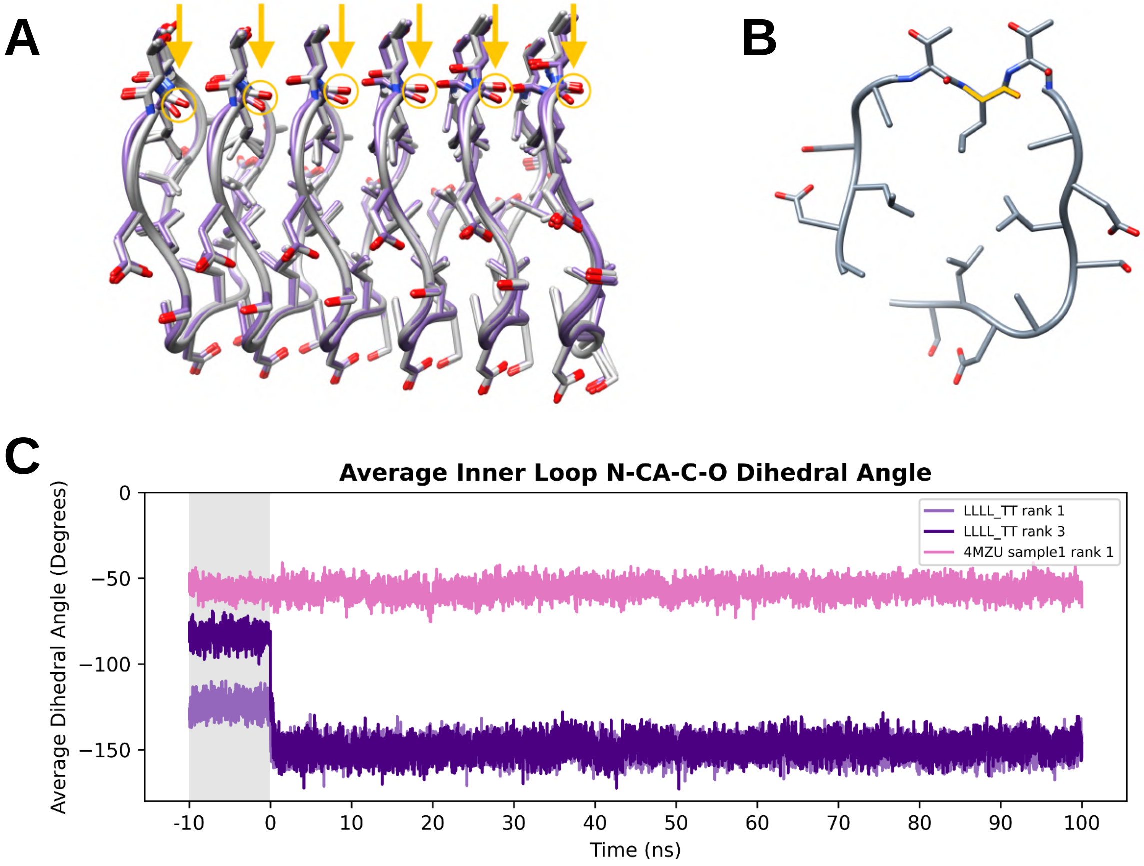


Figure S8: (A) AlphaFold 2 structures of the LLLL_TT model variant, with ranks 1 and 2 colored purple and 3 through 5 colored gray; the arrows highlight the varied X residue carbonyl orientations. (B) A single solenoid loop with yellow lines indicating the X residue N-CA-C-O dihedral angle. (C) Average inner loop N-CA-C-O dihedral angle versus time for 242 K simulations of LLLL_TT rank 1, LLLL_TT rank 3, and 4MZU sample1 rank 1, with the equilibration time shaded gray.

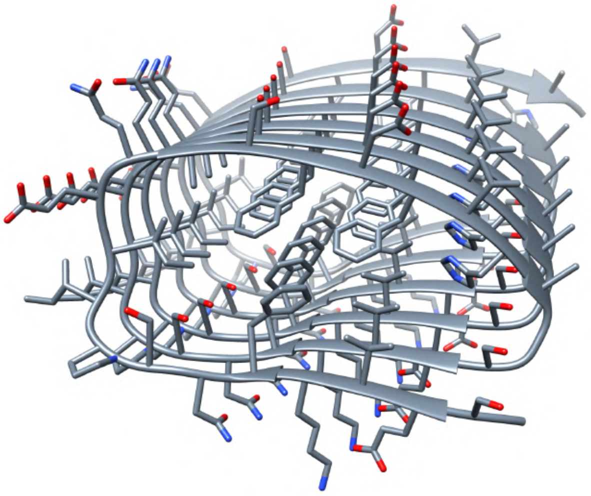

Figure S9: The rank 1 AlphaFold structure of the six-loop construct derived from PDB 4OJ5.

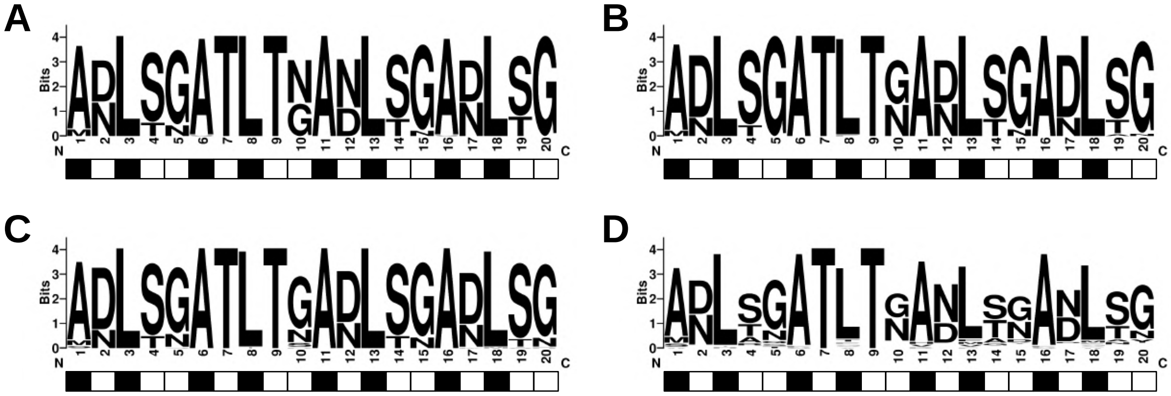


Figure S10: Sequence logos for sequences generated with ProteinMPNN for the rank 1 AlphaFold structure of LLLL_TT with a sampling temperature of (A) 0.1, (B) 0.3, (C) 0.5, and (D) 0.8. Each logo represents 60 loop sequences (six loops for each of 10 full sequences). The grid beneath each logo indicates whether the residue faces inward (black) or outward (white).

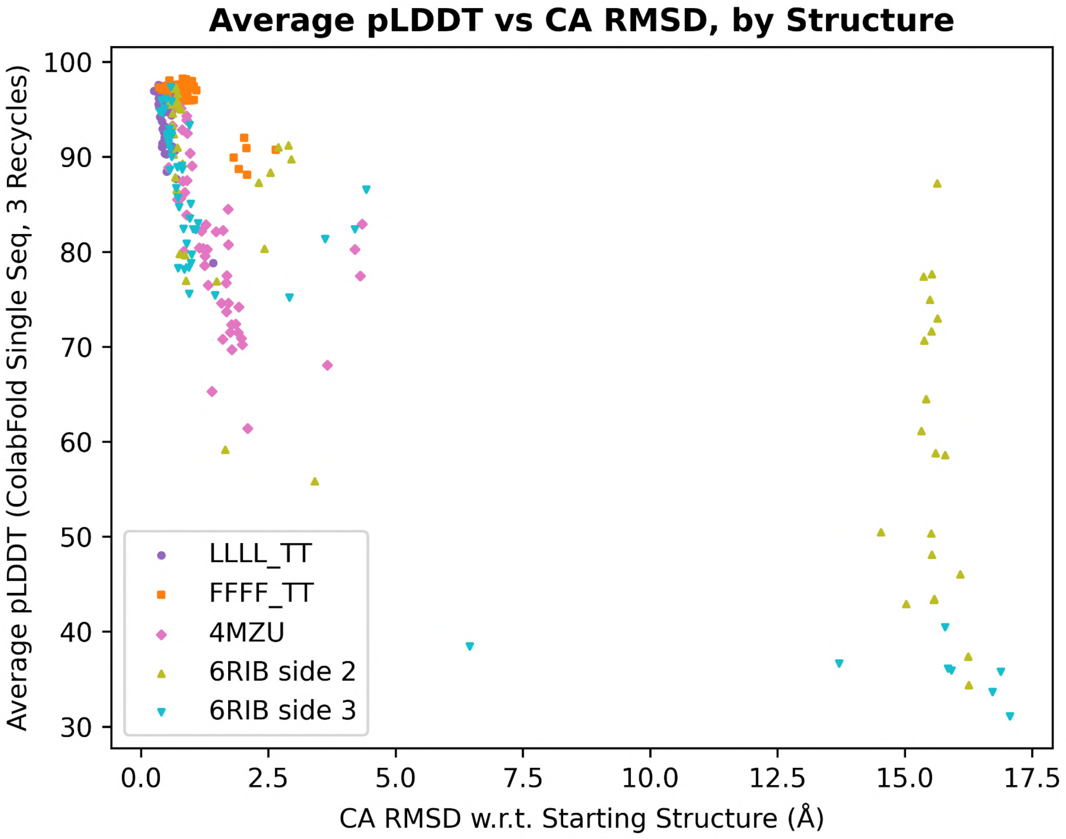


Figure S11: Average pLDDT versus CA RMSD (with respect to the corresponding starting structure provided to ProteinMPNN) for each AlphaFold 2 structure for each sequence generated with ProteinMPNN.

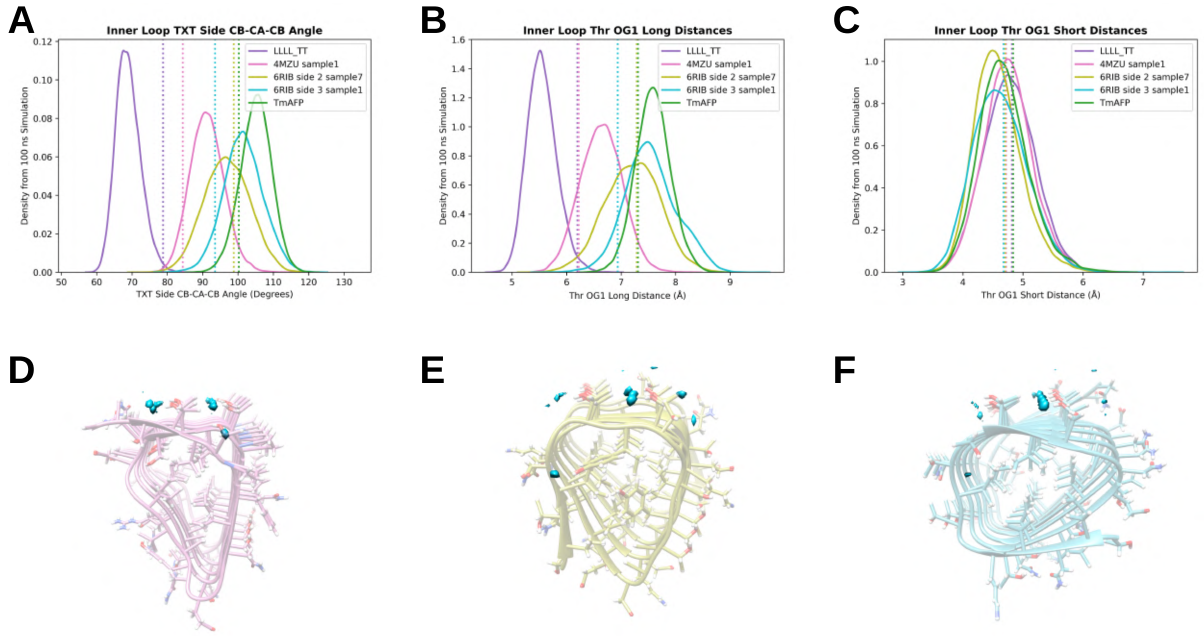


Figure S12: Distributions of inner loop (A) TXT side CB-CA-CB angles, (B) threonine OG1 long distances, and (C) threonine OG1 short distances from 242 K simulations of three non-pentapeptide repeat solenoid sequences from ProteinMPNN, with the distributions from TmAFP and LLLL_TT included for comparison. The OG1 distances are filtered based on the threonines' rotameric states; unfiltered distributions are included in Figure S13 of the Supporting Information. Vertical dashed lines represent the values from the rank 1 AlphaFold structures (or PDB 1EZG chain A for TmAFP). (D) through (F) show the locations of peaks in water oxygen density (threshold of 5%) around the threonine ladders alongside average structures of (D) 4MZU sample1, (E) 6RIB side 2 sample7, and (F) 6RIB side 3 sample1.

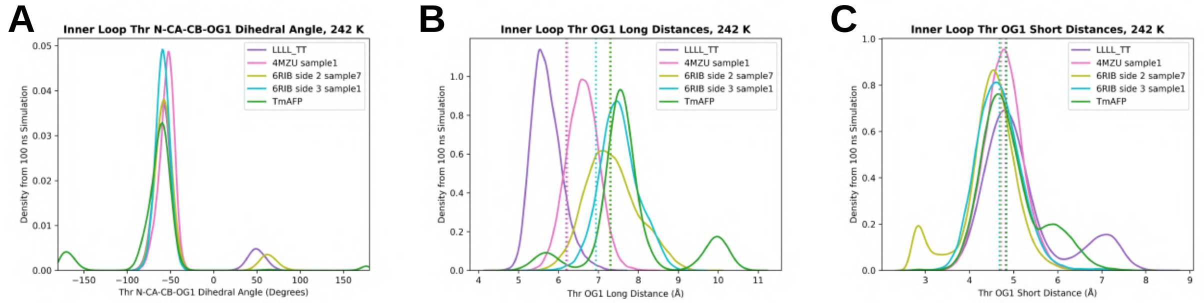


Figure S13: (A) Inner loop threonine N-CA-CB-OG1 dihedral angle distributions, (B) unfiltered threonine OG1 long distance distributions, and (C) unfiltered threonine OG1 short distance distributions at 242 K for the three non-pentapeptide repeat solenoid sequences from ProteinMPNN, with the distributions from TmAFP and LLLL_TT also shown for comparison. In (B) and (C), vertical dashed lines represent the values from the rank 1 AlphaFold structures (or PDB 1EZG chain A for TmAFP).

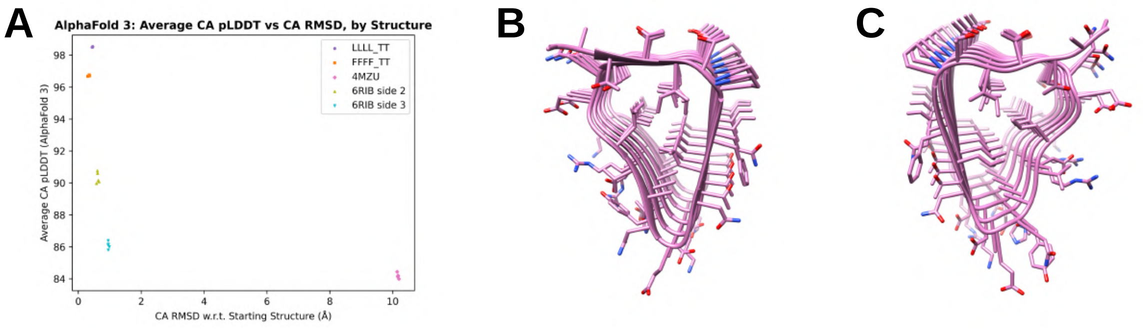


Figure S14: AlphaFold 3 comparison for our selected ProteinMPNN sequences (those in Figure 6E and Table S4). (A) Average alpha-carbon pLDDT versus CA RMSD with respect to the starting structure provided to ProteinMPNN for the five AlphaFold 3 predictions for each sequence. (B) The AlphaFold 2 rank 1 structure of the 4MZU-derived sequence, and (C) the AlphaFold 3 model 0 structure for the same sequence.

**Table S1: Pentapeptide Repeat Variant Loop Sequences (Threonine-Free).**

| Label | Loop Sequence |
|---|---|
| VVVV_DS | ADVSGADVSGADVSGADVSG |
| LLLL_DS | ADLSGADLSGADLSGADLSG |
| FFFF_DS | ADFSGADFSGADFSGADFSG |
| WWWW_DS | ADWSGADWSGADWSGADWSG |
| VLVL_DS | ADVSGADLSGADVSGADLSG |
| VFVF_DS | ADVSGADFSGADVSGADFSG |
| VWVW_DS | ADVSGADWSGADVSGADWSG |
| LVLV_DS | ADLSGADVSGADLSGADVSG |
| LFLF_DS | ADLSGADFSGADLSGADFSG |
| LWLW_DS | ADLSGADWSGADLSGADWSG |
| FVFV_DS | ADFSGADVSGADFSGADVSG |
| FLFL_DS | ADFSGADLSGADFSGADLSG |
| FWFW_DS | ADFSGADWSGADFSGADWSG |
| WVWV_DS | ADWSGADVSGADWSGADVSG |
| WLWL_DS | ADWSGADLSGADWSGADLSG |
| WFWF_DS | ADWSGADFSGADWSGADFSG |
| VLVV_DS | ADVSGADLSGADVSGADVSG |
| VFVV_DS | ADVSGADFSGADVSGADVSG |
| VWVV_DS | ADVSGADWSGADVSGADVSG |
| LVLL_DS | ADLSGADVSGADLSGADLSG |
| LFLL_DS | ADLSGADFSGADLSGADLSG |
| LWLL_DS | ADLSGADWSGADLSGADLSG |
| FVFF_DS | ADFSGADVSGADFSGADFSG |
| FLFF_DS | ADFSGADLSGADFSGADFSG |
| FWFF_DS | ADFSGADWSGADFSGADFSG |
| WVWW_DS | ADWSGADVSGADWSGADWSG |
| WLWW_DS | ADWSGADLSGADWSGADWSG |
| WFWW_DS | ADWSGADFSGADWSGADWSG |
| VVVL_DS | ADVSGADVSGADVSGADLSG |
| VVVF_DS | ADVSGADVSGADVSGADFSG |
| VVVW_DS | ADVSGADVSGADVSGADWSG |
| LLLV_DS | ADLSGADLSGADLSGADVSG |
| LLLF_DS | ADLSGADLSGADLSGADFSG |
| LLLW_DS | ADLSGADLSGADLSGADWSG |
| FFFV_DS | ADFSGADFSGADFSGADVSG |
| FFFL_DS | ADFSGADFSGADFSGADLSG |
| FFFW_DS | ADFSGADFSGADFSGADWSG |
| WWWV_DS | ADWSGADWSGADWSGADVSG |
| WWWL_DS | ADWSGADWSGADWSGADLSG |
| WWWF_DS | ADWSGADWSGADWSGADFSG |

**Table S2: Pentapeptide Repeat Variant Loop Sequences (With Threonines).**

| Label | Loop Sequence |
|---|---|
| VVVV_TT | ADVSGATVTGADVSGADVSG |
| LLLL_TT | ADLSGATLTGADLSGADLSG |
| FFFF_TT | ADFSGATFTGADFSGADFSG |
| WWWW_TT | ADWSGATWTGADWSGADWSG |
| VLVL_TT | ADVSGATLTGADVSGADLSG |
| VFVF_TT | ADVSGATFTGADVSGADFSG |
| VWVW_TT | ADVSGATWTGADVSGADWSG |
| LVLV_TT | ADLSGATVTGADLSGADVSG |
| LFLF_TT | ADLSGATFTGADLSGADFSG |
| LWLW_TT | ADLSGATWTGADLSGADWSG |
| FVFV_TT | ADFSGATVTGADFSGADVSG |
| FLFL_TT | ADFSGATLTGADFSGADLSG |
| FWFW_TT | ADFSGATWTGADFSGADWSG |
| WVWV_TT | ADWSGATVTGADWSGADVSG |
| WLWL_TT | ADWSGATLTGADWSGADLSG |
| WFWF_TT | ADWSGATFTGADWSGADFSG |
| VLVV_TT | ADVSGATLTGADVSGADVSG |
| VFVV_TT | ADVSGATFTGADVSGADVSG |
| VWVV_TT | ADVSGATWTGADVSGADVSG |
| LVLL_TT | ADLSGATVTGADLSGADLSG |
| LFLL_TT | ADLSGATFTGADLSGADLSG |
| LWLL_TT | ADLSGATWTGADLSGADLSG |
| FVFF_TT | ADFSGATVTGADFSGADFSG |
| FLFF_TT | ADFSGATLTGADFSGADFSG |
| FWFF_TT | ADFSGATWTGADFSGADFSG |
| WVWW_TT | ADWSGATVTGADWSGADWSG |
| WLWW_TT | ADWSGATLTGADWSGADWSG |
| WFWW_TT | ADWSGATFTGADWSGADWSG |
| VVVL_TT | ADVSGATVTGADVSGADLSG |
| VVVF_TT | ADVSGATVTGADVSGADFSG |
| VVVW_TT | ADVSGATVTGADVSGADWSG |
| LLLV_TT | ADLSGATLTGADLSGADVSG |
| LLLF_TT | ADLSGATLTGADLSGADFSG |
| LLLW_TT | ADLSGATLTGADLSGADWSG |
| FFFV_TT | ADFSGATFTGADFSGADVSG |
| FFFL_TT | ADFSGATFTGADFSGADLSG |
| FFFW_TT | ADFSGATFTGADFSGADWSG |
| WWWV_TT | ADWSGATWTGADWSGADVSG |
| WWWL_TT | ADWSGATWTGADWSGADLSG |
| WWWF_TT | ADWSGATWTGADWSGADFSG |

**Table S3: LLLL_TT X Residue Carbonyl Data.**

| Predicted Structure | Avg Inner Loop N-CA-C-O Dihedral |
|---|---|
| AF2 Rank 1 | -108.7 |
| AF2 Rank 2 | -108.2 |
| AF2 Rank 3 | -76.4 |
| AF2 Rank 4 | -76.5 |
| AF2 Rank 5 | -76.1 |
| AF3 Model 0 | -151.1 |
| AF3 Model 1 | -151.6 |
| AF3 Model 2 | -151.3 |
| AF3 Model 3 | -151.1 |
| AF3 Model 4 | -150.9 |

**Table S4: Selected ProteinMPNN Sequences.**

| Starting Point | ProteinMPNN Sequence |
|---|---|
| LLLL_TT | MDLSGSTLTGVDLSSSRLSGADLSQATLTI<br>ADLSNADLSGADLKGATLTNADLNDADLSQ<br>ADLSQATLTGADLSNADLSGADITQATLTY<br>ADLSYADLTGADLTGATFTEADVTGADWTG |
| FFFF_TT | HNFSGSTFTGQNFAGTDFSHANFSGATLTK<br>TDFSGANFNNADMSNATFTSTNFDGANFNG<br>ADFTNATFTNTKFNGADFNNAKFNNATFTDT<br>DFNDANFNGADFDGATFTGTKFEGANWTP |
| 4MZU | AATTITPGFLNGENAVLGDNSTLTPGQLTG<br>AYARVGANSTLTPGILTGDTAEVGDSSTLT<br>PGILTGKHARVGANSTLTPGILTGDEAQVG<br>ANSTLTPGILTGRNAVVK |
| 6RIB side 2 | TLTIDGTYTGSYTADEININGTLTAKITAK<br>TIKVKGTVTAEITAEKITIDGTVTGTITAAD<br>IDVNGTVTANITAANLTVAGTMTAEVTTA |
| 6RIB side 3 | TVNVKGTVTTTITANEVIITGTLTGTLTAN<br>SVTINGTVNGTITANIVNITGTLSGTVTAA<br>TVNITGTVTGTITADALTIKGTDTSTVTSN |

### Amber Inputs

```
tLEaP Preparation:

source leaprc.protein.ff14SB

source leaprc.water.tip4pew

SYS = loadPdb "LLLL_TT_unrelaxed_rank_001_alphafold2_ptm_model_5_seed_000.pdb"

solvatebox SYS TIP4PEWBOX 15.0 iso

setBox SYS "centers"

savepdb SYS LLLL_TTr1_solvated_tleap.pdb

saveamberparm SYS LLLL_TTr1.prmtop LLLL_TTr1.rst7

quit
```

```
Minimization:

Minimize with backbone restrained

 &cntrl

  imin=1,

  ntx=1,

  irest=0,

  maxcyc=15000,

  ncyc=5000,

  ntpr=100,

  ntwx=100,

  cut=12.0,

  ntr=1,

  restraintmask='@CA,C,O,N&!:WAT',

  restraint_wt=10.0,

 /
```

Heating:

Heat 0 to 300 K over 50 ps NVT with protein backbone restrained

```
 &cntrl
  imin=0,
  ntx=1,
  irest=0,
  nstlim=25000,
  dt=0.002,
  ntf=2,
  ntc=2,
  tempi=0.0,
  temp0=300.0,
  ntpr=100,
  ntwx=100,
  cut=12.0,
  ntb=1,
  ntp=0,
  ntt=3,
  gamma_ln=2.0,
  nmropt=1,
  ig=-1,
  ntr=1,
  restraintmask='@CA,C,O,N&!:WAT',
  restraint_wt=10.0,
 /
&wt type='TEMP0', istep1=0, istep2=25000, value1=0.0, value2=300.0 /
&wt type='END' /
```

Equilibration:

Equil with weak backbone restraints 10 ns NPT

```
 &cntrl
  imin=0,
  ntx=5,
  irest=1,
  nstlim=5000000,
  dt=0.002,
  ntf=2,
  ntc=2,
  temp0=300.0,
  ntpr=5000,
  ntwx=5000,
  cut=12.0,
  ntb=2,
  ntp=1,
  ntt=3,
  gamma_ln=2.0,
  ig=-1,
  iwrap=1,
  ntr=1,
  restraintmask='@CA,C,O,N&!:WAT',
  restraint_wt=2.0,
 /
```

```
Production:

Production
 &cntrl
  imin=0,
  ntx=5,
  irest=1,
  nstlim=50000000,
  dt=0.002,
  ntf=2,
  ntc=2,
  temp0=300.0,
  ntpr=5000,
  ntwx=5000,
  ntwr=-5000000
  cut=12.0,
  ntb=2,
  ntp=1,
  ntt=3,
  gamma_ln=2.0,
  ig=-1,
  iwrap=1,
 /
```
